# Translatable electrophysiological and behavioral abnormalities in a humanized model of *SYNGAP1*-disorder

**DOI:** 10.1101/2024.08.22.609238

**Authors:** Alex J. Felix, Brandon L. Brown, Nicolas Marotta, Maximilian J. Gessner, Mika Houserova, Icnelia Huerta-Ocampo, Taryn Wilson, Rani Randell, Jennine M. Dawicki-McKenna, Dulce Reinhardt, Keita Uchida, Ian McSalley, Jillian L. McKee, Ingo Helbig, Michael J. Boland, Beverly L. Davidson, Benjamin L. Prosser

## Abstract

Heterozygous variants in *SYNGAP1* and *STXBP1* cause distinct neurodevelopmental disorders due to haploinsufficiency of essential synaptic proteins. As gene targeted approaches to correct these disorders often target non-conserved genomic regions, thus limiting their clinical translation, we generated humanized mouse models wherein the entire *Syngap1* or *Stxbp1* loci were replaced with their human counterparts. *Stxbp1* humanized mice exhibited impaired viability, while *Stxbp1* hybrid mice (*Stxbp1^Hu/+^)* were viable and suitable for evaluating target engagement of human-specific therapeutics. *Syngap1* humanized mice were viable and successfully crossed with *Syngap1* heterozygous mice to produce a *Syngap1* humanized-haploinsufficient model (*Syngap1^Hu/-^*). *Syngap1^Hu/-^* mice displayed haploinsufficient levels of human SYNGAP1, disease-relevant behaviors, and EEG abnormalities including epileptiform activity and generalized slowing. Importantly, parallel analysis in a cohort of patients with *SYNGAP1*-disorder revealed similar electrophysiological signatures. Finally, we showed that human gene-targeted antisense oligonucleotides modulate human *SYNGAP1* expression in *Syngap1^Hu/-^* neurons. Together, we describe new models to support pre-clinical therapeutic development for *SYNGAP1* and *STXBP1* disorders and identify translational biomarkers of *SYNGAP1*-disorder in mice and humans to benchmark therapeutic testing.

## Introduction

*De novo*, heterozygous mutations in Synaptic Ras GTPase Activating Protein 1 (*SYNGAP1*) and Syntaxin Binding Protein 1 (*STXBP1*) cause distinct, rare neurodevelopmental disorders (NDDs) with an incidence of ∼1 in 10,000 and ∼1 in 30,000 births, respectively [1, 2]. Mutations reduce levels of post-synaptic *SYNGAP1* and pre-synaptic *STXBP1*, each of which are required for proper synaptic function and neuroplasticity [3, 4]. *SYNGAP1* and *STXBP1* epileptic encephalopathies are characterized by severe-to-profound intellectual disability, epilepsy, motor dysfunction and autistic features [5–7]. There are no treatments available to alter disease course or that address the genetic cause of these disorders.

Various *Stxbp1* heterozygous knockout mouse models (*Stxbp1^+/−^*) have been generated and validated to study *STXBP1*-disorder *in vivo* [3, 8–10]. Like patients, *Stxbp1^+/−^* mice have haploinsufficient levels of STXBP1 and some phenotypic features associated with STXBP1 loss-of-function, including hyperactivity, impaired cognition, anxiety-like behaviors and motor dysfunction. Mouse models also recapitulate electroencephalographic (EEG) signatures observed in humans, including spike-wave discharges and myoclonic seizures [3, 9]. Similarly, existing mouse models of *SYNGAP1*-disorder (*Syngap1^+/−^* and knock-in models harboring known pathogenic variants) show SYNGAP1 protein haploinsufficiency [11–13], are hyperactive, have memory deficits [11, 14], and show altered EEG power spectra [15].

Several therapeutic modalities to rescue haploinsufficiency disorders are in development and include antisense oligonucleotide (ASO) approaches [16–18], engineered translational activators [19, 20], CRISPR activation and gene editing strategies. These candidate therapies target specific regions of the human gene that are not fully conserved in rodents, precluding *in vivo* testing [8, 12]. This highlights the need for generating animal disease models that incorporate the molecular features of the target human gene to enable *in vivo* testing of human gene-targeted approaches, ultimately accelerating their transition to the clinic.

Towards this goal, we generated and characterized *Syngap1* and *Stxbp1* humanized mouse models wherein the entire *Syngap1* or *Stxbp1* murine loci were replaced with the orthologous human sequences. This included the upstream and downstream regulatory regions, allowing broad utility for different gene-targeted strategies. We next generated a *Syngap1* humanized-disease mouse model that was haploinsufficient. Our data show that *Syngap1^Hu/-^* mice exhibit key disease-related behaviors and distinct EEG abnormalities. *SYNGAP1*-disorder patients that underwent EEG analyses revealed similar signatures including spike-wave discharges and generalized slowing. We also demonstrate that human *SYNGAP1*-targeted approaches can modulate *SYNGAP1* levels in cultured neurons, establishing a platform for pre-clinical development of human-specific gene therapies.

## Materials and Methods

### Generation of *Syngap1* and *Stxbp1* humanized mouse models

The *Syngap1* humanized mouse model is a non-conditional knock-in (KI) model generated by introducing a ∼37.5 kb of gDNA encoding the human *SYNGAP1* and *ZBTB9* genes in place of a ∼34.7 kb of the murine *Syngap1* and *Zbtb9* genes (**Figure 1A**) via Bacterial Artificial Chromosome (BAC) targeting in mouse Embryonic Stem (ES) cells. The entire *Syngap1* locus is humanized (including promoter, proximal enhancers, 5’UTR, coding sequence, intronic regions and the entire 3’UTR), while the *Zbtb9* is only humanized up to the STOP codon (**Figure 1A, magenta box**). This allows for the inclusion of the *SYNGAP1* antisense transcript (*SYNGAP1-AS*) within the humanized region (**Figure 1A, orange box**), to enable testing of therapeutic approaches targeting this element. The final KI human gDNA was flanked with loxP sites. The BAC ID used for the *Syngap1* humanization is RP11-175A4. *Syngap1* humanization may affect the expression of the *Cuta* gene (**Figure 1A, brown boxes**) as *Syngap1* and *Cuta* appear to share a divergently transcribed promoter and approximately half of the region corresponding to this promoter was humanized.

**Figure 1.**
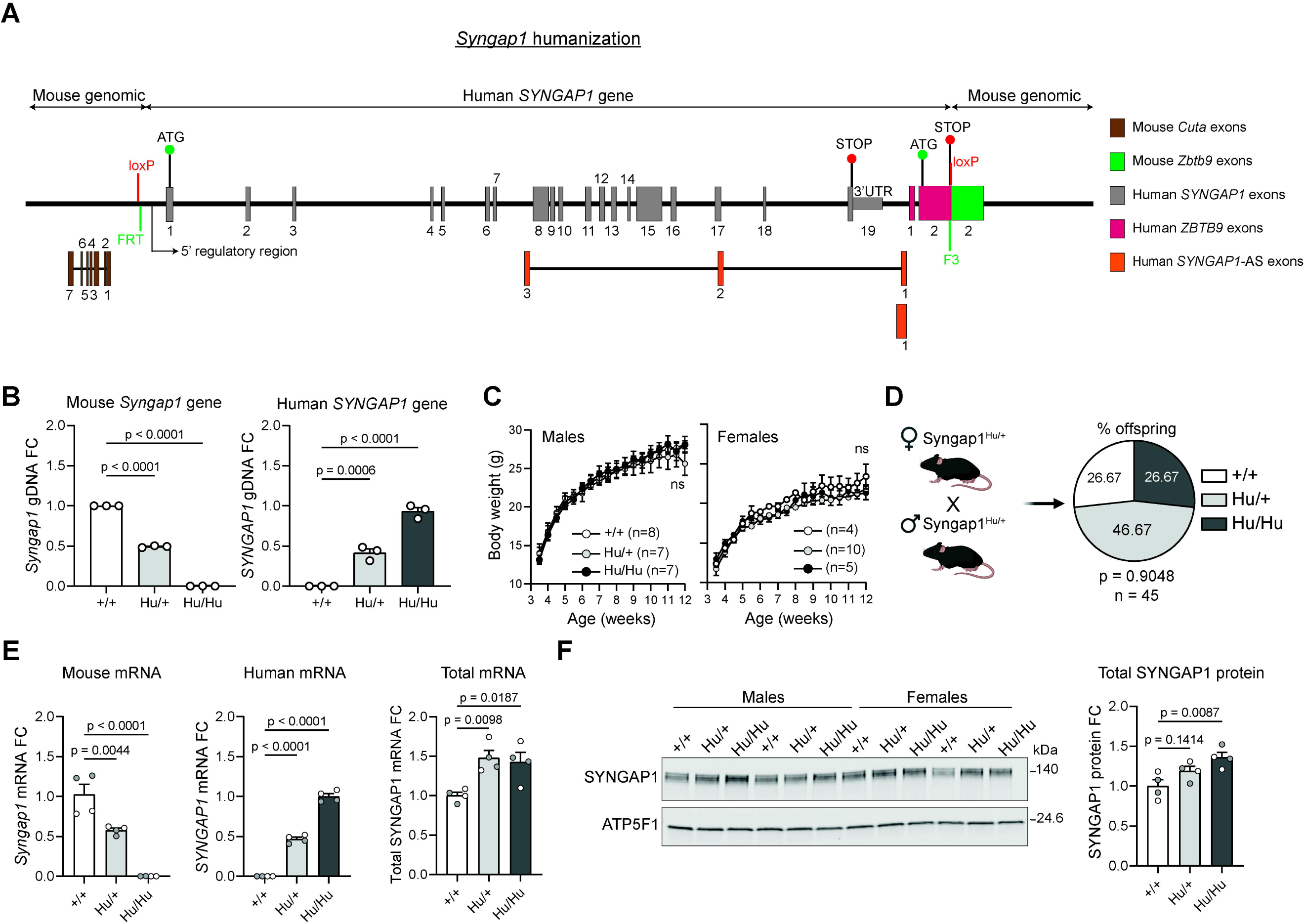
Generation of the *Syngap1* humanized mouse model. (**A**) Cartoon depicting the *Syngap1* locus after humanization. FRT and F3 are two heterotypic recognition sequences used for FLP-mediated neomycin and hygromycin cassette removal. The human *SYNGAP1* transgene is flanked with loxP sites. 5’ regulatory region indicates promoter and proximal enhancers. (**B**) qPCR from gDNA of wild-type (+/+), hybrid (Hu/+) and fully humanized (Hu/Hu) *Syngap1* mice. *Tert* was used as endogenous control. (**C**) Body weights of *Syngap1* model mice. (**D**) Genotypic ratios from the offspring of *Syngap1^Hu/+^*x *Syngap1^Hu/+^* breedings. Created with BioRender. (**E**) RT-qPCR from cerebral cortex tissue of 9-week-old *Syngap1* model mice. *Atp5f1* mRNA was used as endogenous control. (**F**) SYNGAP1 western blot from samples in (**E**). ATP5F1 was used as endogenous control. (**B**, **C**, **E** and **F**) Data are represented as mean values ± SEM. Data points represent independent biological replicates. (**E** and **F**) White and gray data points indicate females and males, respectively. (**B, E** and **F**) One-way ANOVA with Dunnett’s multiple comparison test vs. wild-type (+/+). (**C**) 2-way ANOVA with Dunnett’s multiple comparison test vs. wild-type (+/+) (**D**) Chi-square test (df = 2, *n* = 45, p = 0.9048). *SYNGAP1-AS*, *SYNGAP1* antisense transcript. FC, fold change. ns, non-statistically significant.

The Stxbp1 humanized mouse model is a non-conditional KI model also generated by BAC targeting in mouse ES cells. The BAC replaced the murine genomic region from approximately 5.4 kb upstream of *Stxbp1* exon 1 through to near the end of exon 19 with the human genomic region from approximately 4.0 kb upstream of *STXBP1* exon 1 (Transcript ENST00000373299) through to the end of the final exon of human *STXBP1* (Transcript ENST00000636962). This represents the introduction of ∼89 kb of gDNA encoding the human *STXBP1* (including promoter, proximal enhancers, 5’UTR, coding sequence, intronic regions and the entire 3’UTR) in place of ∼66Kb of the murine *Stxbp1*. The final KI human gDNA was flanked with loxP sites. The humanized region may express the miR-3911 (**Figure 2A, red box**) and lncRNA ENST00000624141 (**Figure 2A, yellow box**), as well as the short 35 residue isoform of PTRH1 (Uniprot: A0A286YER0) (**Figure 2A, orange boxes**). The BAC ID used for the *Stxbp1* humanization is RP11-42D4.

**Figure 2.**
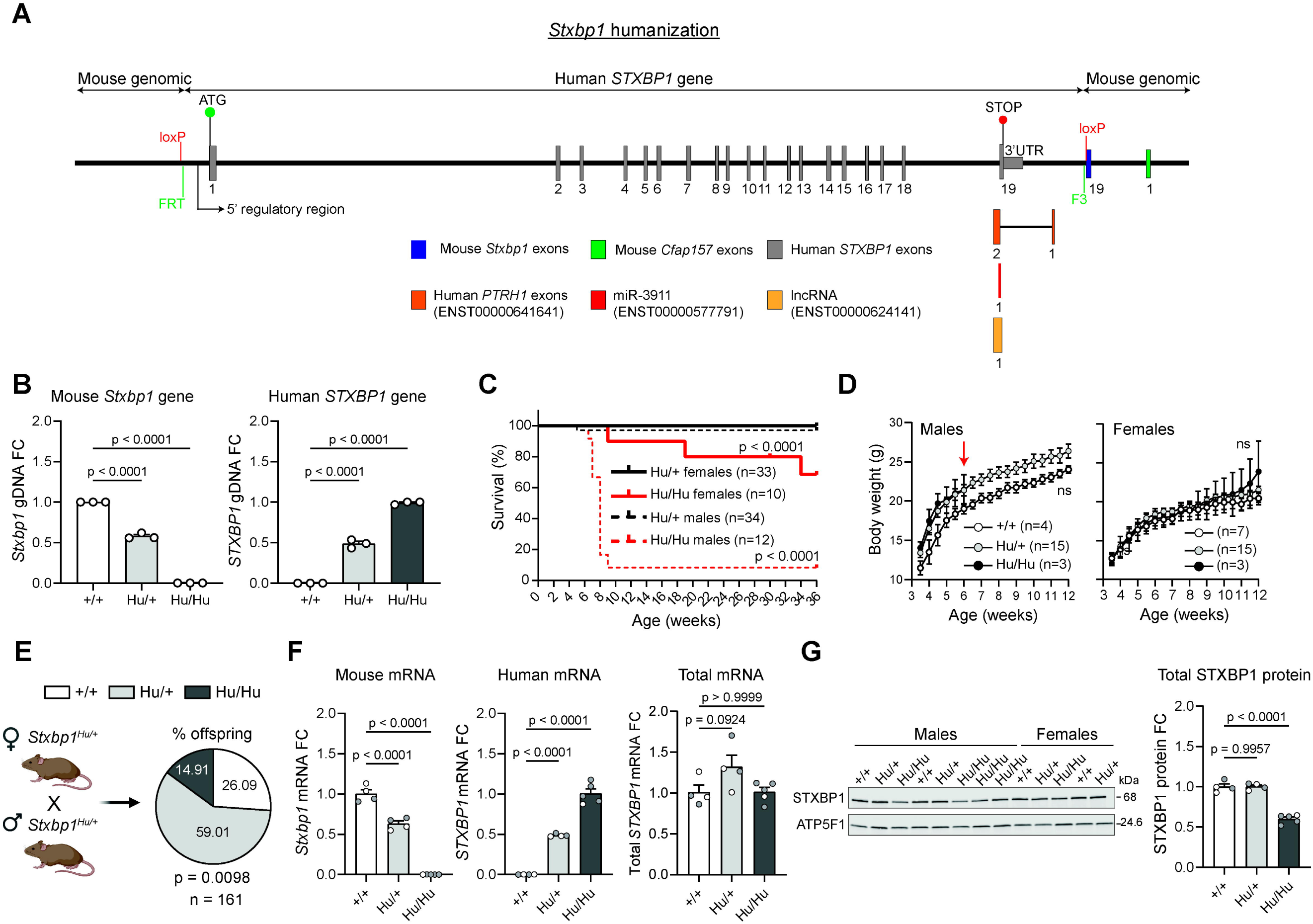
Generation of the *Stxbp1* humanized mouse model. **(A)** Cartoon depicting the *Stxbp1* locus after humanization. FRT and F3 are two heterotypic recognition sequences used for FLP-mediated neomycin and hygromycin cassette removal. The human *STXBP1* transgene is flanked with loxP sites. 5’ regulatory region indicates promoter and proximal enhancers. (**B**) qPCR from gDNA of wild-type (+/+), hybrid (Hu/+) and fully humanized (Hu/Hu) *Stxbp1* model mice. *Tert* was used as endogenous control. (**C**) Kaplan-Meier survival curves of *Stxbp1^Hu/+^* and *Stxbp1^Hu/Hu^* mice. (**D**) Body weights of *Stxbp1* model mice. Red arrow indicates the last data available for male *Stxbp1^Hu/Hu^*. (**E**) Genotypic ratios from the offspring of *Stxbp1^Hu/+^* x *Stxbp1^Hu/+^* breedings. Created with BioRender. (**F**) RT-qPCR from cerebral cortex tissue of 8-week-old *Stxbp1* model mice. *Atp5f1* mRNA was used as endogenous control. (**G**) STXBP1 western blot from samples in (**F**). ATP5F1 was used as endogenous control. (**B**, **D**, **F** and **G**) Data are represented as mean values ± SEM. Data points represent independent biological replicates. (**F** and **G**) White and gray data points indicate females and males, respectively. (**B, F** and **G**) One-way ANOVA with Dunnett’s multiple comparison test vs. wild-type (+/+). (**C**) Mantel-Cox test. (**D**) 2-way ANOVA with Dunnett’s multiple comparison test vs. wild-type (+/+). (**E**) Chi-square test (df = 2, *n* = 161). FC, fold change. ns, non-statistically significant.

All mouse gene engineering steps were performed by Ozgene. The BAC constructs for *Syngap1* and *Stxbp1* humanization were constructed by a third-party who performed quality control via restriction digestion and Pulse-Field Gel Electrophoresis. Inserted cassettes and additional modified parts of the BACs were confirmed by PCR and sequencing. Ozgene also performed independent quality controls by sequencing or qPCR of key regions such as loxP sites, neomycin and hygromycin cassettes, junctions, and presence of *SYNGAP1* and *STXBP1* transgenes. The BACs were electroporated into C57BL/6 ES cells and qPCR assays were carried out to confirm correct targeting as well as presence of the selection cassettes and the corresponding inserts. Gene-targeted ES cell clones were injected into goGermline blastocysts to produce goGermline chimeras followed by F1 heterozygous targeted mice (Hu/+) in which the selection cassettes (neomycin and hygromycin) were removed by mating the chimeras to a ubiquitous Flp line.

The resulting *Syngap1* and *Stxbp1* humanized mouse models are made available through JAX under MMRRC_069939 and MMRRC_071410, respectively.

### Generation of S*yngap1* and *Stxbp1* humanized-disease mouse models

B6;129-*Syngap1^tm1Rlh^*/J mice were obtained from The Jackson Laboratory (JAX #008890). To generate the *Syngap1* humanized-disease mouse model, male mice B6;129-Syngap1^tm1Rlh^/J (*Syngap1^+/−^*) were bred with *Syngap1^Hu/Hu^* females to generate *Syngap1^Hu/-^* and wild-type *Syngap1^Hu/+^* littermates.

B6;129S-*Stxbp1^tm1Sud^*/J mice were obtained from The Jackson Laboratory (JAX #006381), and male mice B6;129S-*Stxbp1^tm1Sud^*/J (*Stxbp1^+/−^*) were crossed with either *Stxbp1^Hu/+^* or *Stxbp1^Hu/Hu^* females to try to generate *Stxbp1^Hu/-^* and wild-type *Stxbp1^Hu/+^*littermates.

Genotyping for B6;129-*Syngap1^tm1Rlh^*/J and B6;129S-*Stxbp1^tm1Sud^*/J mice was performed by Transnetyx using real-time PCR or in-house using the primer-probe sets in **Supplementary Table 1**.

### Animals

The *Syngap1* humanized mouse model was maintained on a pure C57BL/6J genetic background, while B6;129-*Syngap1^tm1Rlh^*/J mice were maintained on a mixed background of 129S1/SvImJ and C57BL/6J. The *Syngap1* humanized-disease mouse model was on a mixed background of 129S1/SvImJ and C57BL/6J. The *Stxbp1* humanized model was maintained on a mixed genetic background of C57BL/6J outcrossed one generation to the BALB/c strain. B6;129S-*Stxbp1^tm1Sud^*/J mice were maintained in a pure C57BL/6J background. *Stxbp1* humanized-disease mice were on a mixed background of C57BL/6J and BALB/c strain.

Both male and female mice were used in this study. Mice were maintained on a 12:12-h light:dark cycle with *ad libitum* access to food and water and were weaned at 21 days. For *Syngap1* and *Stxbp1* humanized mouse models, body weight measurements were obtained twice per week until animals reached 12 weeks of age. For the *Syngap1* humanized-disease mouse model, body weights were recorded weekly until mice reached 15 weeks of age. Survivability was assessed up to 36 weeks of age.

Both male and female mice were generated at the Children’s Hospital of Philadelphia (CHOP) for behavioral phenotyping. Genetic background of these mice (*Syngap1^Hu/-^* and *Syngap1^Hu/+^)* were confirmed by miniMUGA testing of 1 mouse per litter and showed a genetic background of ∼85% C57BL/6J.

### Copy number variation assay

Ear samples or tail snips from mice were collected in 1.5 mL tubes for genotyping. Genomic DNA (gDNA) extraction was performed by adding 100 µL of DirectPCR Lysis Reagent (Viagen Biotech #402-E) supplemented with proteinase K (Viagen Biotech #505-PKP) and incubating the samples at 56 °C overnight. Proteinase K was then inactivated at 86 °C for 45 min and samples were centrifuged at 8,000 x g for 1 min. The supernatant containing gDNA was directly used for qPCR-based genotyping or stored at 4 °C.

The copy number variation assay with qPCR was prepared by mixing the following reagents: 1 µL of crude gDNA, 1X PrimeTime Gene Expression Master Mix (IDT #1055772), 1X primers/probe mix and nuclease-free water to a final volume of 10 µL. Three technical replicates were performed for each sample. qPCR was carried out on a QuantStudio 3 Real-Time PCR System (ThermoFisher) with a passive reference of ROX using the following cycling conditions: 95 °C for 3 min for 1 cycle, 95 °C for 15 s and 62 °C for 1 min for 40 cycles. ΔCt was calculated by subtracting the average Ct of the reference gene from the average Ct of the gene of interest for each sample. ΔΔCt values were obtained by subtracting the average ΔCt value of the control sample from the ΔCt of the test samples and then converted into 2^−ΔΔCt^ to obtain the fold change of gene expression. Mouse *Tert* was used as endogenous control. In all reactions, samples from WT animals (+/+) were included to determine a reference Ct value corresponding to the presence of two copies of the mouse allele.

### RNA isolation and RT-qPCR

Isolated brain tissue (∼1/4 of a cortex for *Stxbp1* and *Syngap1* humanized mouse models or hemi-brain for *Syngap1* humanized-disease mice) was mixed with 1 mL of TRIzol reagent (Invitrogen #15596018) and a 5 mm stainless steel bead (Qiagen #69989) in an RNase-free microcentrifuge tube. Brain tissue was then homogenized in a TissueLyser LT homogenizer (Qiagen #85600) for 5 min at 50 Hz. The homogenate was centrifuge at 12,000 x g for 5 min at 4 °C and seated for an additional 5 min to precipitate insoluble debris. The supernatant was transferred to a new tube, mixed with 200 µL of chloroform (Acros Organics #190764), shaken vigorously and centrifuged at 12,000 x g for 15 min at 4 °C. Aqueous phase was transferred to a new tube containing 500 µL of ice-cold isopropanol (Sigma-Aldrich #190764) followed by incubation for 10 min on ice and centrifugation at 12,000 x g for 10 min at 4 °C to precipitate RNA. Supernatant was discarded and 1 mL of ice-cold 75% ethanol (Decon laboratories #2701) was added to wash the pellet followed by centrifugation at 7,500 x g for 5 min at 4 °C. Supernatant was again discarded and RNA pellet was air-dried for 20 min. RNA was resuspended in 100 µL of RNase-free water and allowed to reconstitute for 10 min at 56 °C. To ensure RNA integrity for downstream applications, the resuspended RNA was purified using the Quick RNA Miniprep kit (Zymo #R1055) following manufacturer’s instructions. Purified RNA was resuspended in 50 µL of RNase-free water. RNA concentration was determined by measuring OD_260_ nm absorbance in a Synergy HTX reader (Biotek).

cDNA synthesis was performed using the SuperScript IV First-Strand Synthesis System with ezDNase Enzyme (ThermoScientific #18091300) using random hexamer primers according to manufacturer’s instructions. The ezDNase treatment step was performed for all conditions.

Probe-based qPCR was prepared by mixing the following reagents: 1 µL of cDNA, 1X PrimeTime Gene Expression Master Mix (IDT #1055772), 1X primers/probe mix and nuclease-free water to a final volume of 10 µL. Three technical replicates were performed for each sample. qPCR was carried out on a QuantStudio 3 Real-Time PCR System (ThermoFisher) with a passive reference of ROX using the following cycling conditions: 95 °C for 3 min for 1 cycle, 95 °C for 5 s and 60 °C for 30 s for 40 cycles. ΔCt was calculated by subtracting the average Ct of the reference gene from the average Ct of the gene of interest for each sample. ΔΔCt values were obtained by subtracting the average ΔCt value of control samples from the ΔCt of the test samples and then converted into 2^−ΔΔCt^ to obtain the fold change of gene expression.

All qPCR primer-probe sets sequences can be found in **Supplementary Table 1**

### Protein isolation and Western blot

For characterization of the *Syngap1* and *Stxbp1* humanized mouse models, isolated cortices (∼1/4 of cortex per animal) were mixed with 500 µL of 1.5x Laemmli buffer [15% glycerol (Amresco #M152), 3% SDS (Sigma #L5750), 3.75 mM EDTA (Bio-Rad #1610729) and 75 mM Tris, pH 7.5 (Invitrogen #15567027)] in a 2 mL sample tube (Qiagen #990381), and a 5 mm stainless steel bead (Qiagen #69989) was added. Tissue was then homogenized in a TissueLyser LT homogenizer (Qiagen #85600) for 5 min at 50 Hz followed by incubation at 95 °C during 10 min. Protein lysates were briefly spun down and the supernatants were transferred to a new microcentrifuge tube. For characterization of the *Syngap1* humanized-disease mouse model, hemi-brains were mixed with 750 µL of 1X RIPA buffer (Cell Signaling Technology #9806S) supplemented with phosphatase (PhosSTOP, Roche #4906845001) and protease (cOmplete, Roche #11836170001) inhibitor cocktails and homogenized as described above.

Protein extracts were quantified using the Pierce 660nm Protein Assay Kit (Thermo #22662) supplemented with Ionic Detergent Compatibility Reagent (Thermo #22663) or the Pierce BCA Protein Assay Kit (Thermo #23225) in a 96-well plate, according to manufacturer’s protocol. Samples were diluted to the same final concentration, mixed with 1x Orange G dye (Sigma #O3756) containing 10% β-mercaptoethanol (Sigma #M3148) and incubated 10 min at 100 °C before loading. Precast 4-15% TGX protein gels (Bio-Rad #4561086) were loaded with 10-20 µg of total protein lysate and run for 1h at 135V. Proteins were transferred to 0.45 µm nitrocellulose membrane (Bio-Rad #1704271) with a Trans-Blot Turbo Transfer system (Bio-Rad) using the pre-determined high molecular weight transfer protocol (10 min, 2.5 A constant). Blocking of the membrane was performed using Intercept (TBS) Blocking Buffer (LI-COR #927-60001) for at least 1h at room temperature. Incubation with primary antibodies (all diluted 1:1000 in blocking buffer containing 0.1% Tween-20) was carried out overnight at 4 °C. Rabbit anti-SYNGAP1 (Cell Signaling Technology #5539S), rabbit anti-STXBP1 (Munc18-1, Cell Signaling Technology #13414S), mouse anti-phospho-ERK1/2 (Cell Signaling Technology #9106), rabbit anti-ERK1/2 (Cell Signaling Technology #9102) and mouse anti-ATP5F1 (Abcam #ab117991) were used. Membrane was then rinsed with 1x Tris-buffered saline with 0.1% Tween (TBST) 4 times for 5 min. Incubation with secondary antibodies (diluted 1:10,000 in blocking buffer containing 0.1% Tween-20) was performed at room temperature for 1h. For STXBP1 and ERK blots, IRDye 680RD anti-rabbit (LI-COR #926-68073) and IRDye 800CW anti-mouse (LI-COR #926-32212) were used. For SYNGAP1 blots, IRDye 800CW anti-rabbit (LI-COR #926-32213) and IRDye 680RD anti-mouse (LI-COR #926-68072) were used. Membrane was rinsed again with 1x TBST 4 times for 5 min and imaged on Odyssey Imager (LI-COR) using a resolution of 169 µm.

Western blot quantifications were normalized to ATP5F1 according to LI-COR’s Housekeeping Protein Normalization Protocol. A standard curve was included in each blot to ensure assay linearity.

### Behavioral assessments

Animals were transported in their home cages to testing room(s), habituated for 1-2 hours, and all testing apparatuses were thoroughly disinfected before testing and between animals. At 4-weeks and 30-weeks of age, animals were weighed and underwent open field, elevated plus maze, and horizontal ladder testing on nonconsecutive days. For open field testing, animals were individually placed in well-lit, 10.75 × 10.75 x 8 inch plexiglass chambers which were housed in a sound attenuating cubicles (Med Associates Inc., ENV-51050-A). Animals were tracked for 30 minutes using Activity Monitor (Med Associates Inc, Version 6.02) to assess; overall activity levels via total distance traveled, stereotypy (abnormal repetitive behaviors) via number of stereotypic events, and anxiety-like behaviors via time spent in the center of the testing chamber. As an additional test for anxiety-like and exploratory behaviors, elevated plus maze was performed. Animals were individually placed in the center of the maze, continuously recorded for 10 minutes, and tracked using ANY-Maze (version 6.0). Center of mass was used to quantify the time in open arms and number of open arm entries.

Horizontal ladder testing was used to assess locomotor coordination and precision. A 64 cm long ladder with rungs spaced 2 cm apart was elevated 17 cm above the testing table and a hollow black escape box placed at one end. Animals were placed at the starting (open) end of the ladder and following a successful pass were kept in the dark escape box for 1-2 minutes before starting the next pass. Following a “training” pass to familiarize animals with the apparatus and escape box, 5 successful passes were recorded. Passes were considered successful if the animal traversed the ladder without turning around or pausing for >2 seconds. If unsuccessful, animals were replaced at the start. The same blinded observer scored all videos and foot slips were counted if the animal’s paw broke the plane of the ladder rungs.

To assess spatial learning and memory, animals underwent Barnes maze testing consisting of 4 training days and 1 test day, at 5-weeks and 31-weeks of age. The Barnes Maze table (San Diego Instruments; 36 inch diameter table; 20, 2 inch holes) was positioned in the center of well-lit room with black walls, discrete visual cues (large white shapes) on each of the 4 walls, and a video camera positioned directly above the table. Videos were recorded and analyzed using ANY-Maze (version 7.3). On training days all holes were occluded except the “target hole” which was unblocked, allowing mice to enter a dark escape box. The target hole remained in the same position for each animal during training days and was changed every 4 animals to minimize the influence of any unknown cues or preferences. Training days consisted of two non-consecutive trials during which animals were placed in the middle of the table and covered with a cardboard box (15 cm x 15 cm) for 1 minute to habituate. The box was removed and the animal recorded for 150 seconds. If the animal did not successfully enter the target hole after 150 seconds, it was gently guided into the target hole and remained there for 60 seconds. On the test day, the target hole was occluded, and animals were recorded for a single 150 second trial.

### Rodent EEG acquisition and processing

For bilaterally recordings of the barrel cortices, auditory cortices, visual cortices, motor cortices, and CA1 of the hippocampi, recording electrodes were constructed and implanted as previously described [21]. 4 *Syngap1^Hu/+^* and 7 *Syngap1^Hu/-^* mice were implanted between 6-8 weeks old. 48 hours after implantation animals were continuously recorded for up to 72 hours with an acquisition rate of 20,000Hz.

EEG analyses were performed in MATLAB using a custom built analysis program (https://github.com/DavidsonLabCHOP/Felix-Brown_EEGcode_2025). Recordings were down sampled to 2,500Hz and segmented into 30-minute epochs. Recording quality was determined by evaluating root-mean-square error and overall skewness. Epochs with a root-mean-square error of less than 30µV or greater than 200µV or skewness greater than 0.4 were removed from further analyses. Spike-wave discharges were defined as having amplitudes 2.5 times larger than baseline root-mean-square amplitude, a frequency of 6-10 Hz, and lasting at least 1 second [3, 22, 23]. For quantification, spike wave discharges were only counted if occurring in at least 4 leads at once.

For power spectra analysis, filtered recordings were divided into 5-second epochs and epochs containing artifacts (defined as a root-mean-square error z-score ≥3) were removed. Fast Fourier transform was performed, and power spectra was quantified of each major frequency band (delta 0-4 Hz, 4-8 theta Hz, alpha 8-13 Hz, beta 13-25 Hz, and gamma 25-50 Hz), and power for each band across all epochs were averaged. The area under the curve for each frequency band was calculated, and bilateral recordings from the same regions were averaged for each animal. To calculate alpha-delta and alpha-theta ratios, the mean total power for each major frequency band per animal was used. For spike detection, spikes were defined as having a voltage deflection greater than 5 standard deviations above the mean amplitude, occurring at least 200ms apart, and having a width of 50 - 200ms.

### Participant EEG acquisition and analysis

We collected EEGs from retrospective routine exams recorded within the CHOP care network. *SYNGAP1* patients were identified by an existing clinical diagnosis within their Electronic Health Record (EHR). Putative control patients were first selected as those having only normal EEGs on record. We next constructed a list of unique ICD9/10 codes, and a panel of three epileptologists created exclusionary criteria for codes anticipated to affect cerebral function or influence EEG readings. Putative control patients with at least one year of patient history post-EEG and no intersection with the exclusion codes were retained as controls. Within both *SYNGAP1* and control cohorts, any EEG recorded during an emergency visit was omitted.

We included all clinical EEGs meeting the above criteria; in the event of extended outpatient, inpatient long-term monitoring (LTM), or long-term ambulatory home monitoring EEGs, recordings were truncated to the first 4 hours. All EEGs were collected using the standard 10-20 system with the machine reference placed between electrodes ‘Cz’ and ‘Fz’. For each recording, we applied a 60 Hz harmonic filter and 95 Hz infinite impulse response (IIR) filter for anti-aliasing. To normalize between acquisition technologies, all EEG recordings were down sampled to 200 Hz. Artifact removal (e.g. blinks, cardiac noise) was performed through an automated independent component analysis (ICA) pipeline via MNE-ICALabel [24, 25]. We then applied a 2nd order bandpass filter from 0.5-70 Hz and a Laplacian montage. Through manual review we constructed an annotation handling ruleset to avoid events conflicting with resting state (e.g. sleep, seizures). After this preprocessing, we screened recordings for uninterrupted four-second epochs, rejecting any recordings with less than 15 qualifying epochs.

To calculate bandpowers, we applied Welch’s method, at 1 Hz intervals, for each EEG electrode across all epochs, reporting the median power for each electrode at each frequency. Relative power was derived in powerbands described as delta (1-4Hz), theta (4-8Hz), alpha (8-13Hz), and beta (13-30Hz). For reporting powerbands globally and regionally, the median relative power was found between all associated electrodes (where frontal: (‘Fp1’, ‘Fp2’, ‘F3’, ‘F4’, ‘Fz’), temporal: (‘T3’, ‘T4’, ‘T5’, ‘T6’, ‘F7’, ‘F8’,), occipital: (‘O1’, ‘O2’), parietal: (‘P3’, ‘P4’, ‘Pz’), and central: (‘C3’, ‘C4’, ‘Cz’)).

Using EEG reports archived in the EHR for *SYNGAP1* recordings, we extracted clinician annotations regarding the occurrence of spike-wave discharges. These were ordinally characterized as ‘None’, ‘Occasional’, or ‘Frequent’.”

For comparison of power spectra and calculations of alpha-delta and alpha-theta ratios, area under the curve for each major frequency band (Delta 1-3Hz, Theta 4-7Hz, Alpha 8-12Hz, Beta 13-29Hz, and Gamma 20-70Hz) were calculated per brain region. Leads were grouped into the following brain regions: temporal (leads T3, T4, T5, T6, F7, and F8), parietal (leads Pz, P3, and P4), central (leads Cz, C3, and C4), frontal (leads Fp1, Fp2, Fz, F3, and F4), and occipital (leads O1 and O2).

### Primary cortical neuron isolations and ASO treatments

Primary cortical neurons were isolated at embryonic day 16-18 from pregnant *Syngap1^Hu/Hu^* females crossed with *Syngap1^+/−^*males. Cortical tissue was isolated from individual embryos in 1x HBSS (Gibco #14170-112) supplemented with 1% Penicillin/Streptomycin (Gibco #15140122), 1% HEPES (Gibco #15630080) and 1% Sodium Pyruvate (Gibco #11360070) and cut into small pieces (∼2 mm). Cortices were stored on ice and pooled based on genotyping. In brief, DNA from embryos was extracted using the Kapa Mouse genotyping kit (Roche #41106300) and diluted 1:100 for qPCR using SYBR Green (Bio-RAD #1725271) with recommended primers from Jax (strain #008890) to detect the presence of the WT or null *Syngap1* allele. Cortices were digested via incubation in 20 U/mL Papain plus DNase (Worthington Biochemical Corporation, #LK003178 and #LK003172) for 15-20 minutes at 37°C in 1X HBSS with supplements. Digestion was halted by the addition of 10% Heat inactivated-FBS (Corning #MT35-010-CV) and washed once in Neurobasal media (Gibco #21103049) and then Neurobasal supplemented with 1% B27 Plus (Gibco #A3582801), 1% GlutaMax (Gibco #35050061), and 1% Penicillin/Streptomycin. Cortices were manually dissociated into single cells by trituration with a 5 ml Serological pipette 3-4 times followed by 25-30 times with a p1000 pipette tip in 2-3 mL of supplemented Neurobasal media. Live cells were counted with trypan blue on a hematocytometer. Cells (200,000) were seeded in 24-well plates (Corning #353226). Plates were coated with 50 µg/mL of poly-D-lysine (Sigma-Aldrich #P1149) in 0.1 M Borate Buffer, pH 8.5 in a sterile environment over night at room temperature. The next day, plates were washed at least 3 times in sterile H_2_O and incubated in a humidified environment at 37°C in 5% CO2 in supplemented Neurobasal media with 5 µg/mL of laminin (Gibco #23017015). After seeding, cells were incubated in a humidified environment at 37°C in 5% CO_2_, and the following day media was changed. Subsequently, 50% media changes were performed once per week until day in vitro (DIV) 10, and twice a week thereafter.

ASO treatments were performed via gymnotic delivery of the ASO in DIV5-7 *Syngap1^Hu/-^* primary neurons for 7 days. ASOs were purchased from IDT and sequences can be found in **Supplementary Table 2**.

### Statistical analyses

Measurements were taken from distinct samples. The number of samples is stated explicitly in the figure and represented as individual data points for bar graphs. Kaplan–Meier survival curves were used to represent the survivability of the different model mice and the Mantel–Cox (Log-rank) test was used to statistically compare the overall survival between groups. Statistical significance was defined as p < 0.05. Data plots and statistical analyses were performed in GraphPad Prism 10.1 software. Linear mixed model was performed in R (Version 2024.12.1+563). Individual statistical tests applied to each data set are given in the respective figure legends. Data are represented as mean values ± standard error of the mean (SEM).

### Study approval

Mouse breeding and procedures were performed at the University of Pennsylvania Perelman School of Medicine animal facility or at CHOP in accordance with the standards set forth by the University of Pennsylvania Institutional Animal Care and Use Committee or CHOP’s Institutional Animal Care and Use Committee and the Guide for the Care and Use of Laboratory Animals published by the US National Institutes of Health under protocol #807524 and #1358.

All human EEG recordings were obtained for clinical indications at CHOP and analyzed retrospectively. The CHOP Institutional Review Board (IRB) has waived the requirement for consent under IRB protocol 20-017641.

### Data availability

All the data that support the findings of this study are provided in the article and its supplementary information files.

## Results

### Generation of a knock-in *Syngap1* humanized mouse model

The *Syngap1* humanized mouse is a non-conditional KI model in which the mouse *Syngap1* locus was replaced with the human *SYNGAP1* gene (**Figure 1A**). We generated hybrid (*Syngap1^Hu/+^*), fully humanized (*Syngap1^Hu/Hu^*) and wild-type littermate (*Syngap1^+/+^*) animals for which we confirmed the presence of 1, 2 and 0 copies of the human *SYNGAP1* transgene in the targeted mouse locus using a CNV qPCR assay (**Figure 1B**). Grossly, hybrid and humanized *Syngap1* mice were indistinguishable from pure wild-type littermates.

A 36-week-long survivability study of male and female mice with hybrid, fully humanized and wild-type genotypes demonstrated no change in body weights, growth rates, or viability, indicating that both mono- and bi-allelic humanization of the *Syngap1* locus are well tolerated (**Figure 1C**). Genotypic ratios calculated from the offspring of *Syngap1* hybrid matings (*Syngap1^Hu/+^* x *Syngap1^Hu/+^*), which should result in 50% of *Syngap1^Hu/+^*, 25% of *Syngap1^Hu/Hu^*and 25% of *Syngap1^+/+^* mice, indicated normal Mendelian inheritance (**Figure 1D**).

The *Syngap1* humanized mouse model was characterized at the RNA and protein level with cerebral cortex samples. As expected, *Syngap1^Hu/Hu^*mice expressed only human *SYNGAP1* transcript, while *Syngap1^Hu/+^* showed a ∼50% reduction in human *SYNGAP1* mRNA levels relative to *Syngap1^Hu/Hu^* (**Figure 1E**, middle panel). Human *SYNGAP1* expression inversely correlated with mouse *Syngap1* expression, confirming successful humanization of the locus (**Figure 1E**, left panel). Using a cross-reactive qPCR assay to detect both mouse and human *SYNGAP1* transcripts, there was a 1.4x increase in total *SYNGAP1* mRNA in both *Syngap1^Hu/+^*and *Syngap1^Hu/Hu^* mice (**Figure 1E**, right panel), indicating that *SYNGAP1* is efficiently transcribed following humanization of its genomic locus. Western blotting using a SYNGAP1 antibody raised against a fully conserved region of the protein surrounding Arg1070 (mouse and human SYNGAP1 are 99% conserved at the amino acid level) revealed that *Syngap1^Hu/Hu^*had ∼35% increased total SYNGAP1 protein levels relative to wild-type littermates, with a subtler potential increase with mono-allelic humanization (**Figure 1F**). These data point towards modestly enhanced transcription of the human *SYNGAP1* gene and/or increased stability of the human *SYNGAP1* mRNA in humanized mice.

### Generation of a knock-in *Stxbp1* humanized mouse model

Similarly, the *Stxbp1* humanized mouse model was generated by replacing the mouse *Stxbp1* locus with the human *STXBP1* gene (**Figure 2A**). We confirmed the presence of the expected copies of human *STXBP1* transgene in *Stxbp1^Hu/+^* (1 copy), *Stxbp1^Hu/Hu^* (2 copies) and *Stxbp1^+/+^* mice (0 copies) (**Figure 2B**). In terms of physical appearance, *Stxbp1* model mice occasionally presented abnormal tail morphologies (i.e. kinked tails).

Bi-allelic humanization of the *Stxbp1* gene significantly reduced viability. Male *Stxbp1^Hu/Hu^* presented an abrupt ∼90% mortality rate around 9 weeks of age, while *Stxbp1^Hu/Hu^* females showed a milder, more stepwise mortality phenotype with a ∼30% mortality at 36 weeks (**Figure 2C**). However, these observations were not recapitulated in either male or female *Stxbp1^Hu/+^* mice, suggesting that mono-allelic but not bi-allelic humanization is well tolerated (**Figure 2C**). Male *Stxbp1^Hu/Hu^* mice body weights were not significantly different from hybrid or wild-type littermates, discarding growth impairments as a potential cause of death (**Figure 2D**). When evaluating the genotypic distribution from the offspring of *Stxbp1* hybrid matings (*Stxbp1^Hu/+^* x *Stxbp1^Hu/+^*), we observed a significant underrepresentation in the number of weaned *Stxbp1^Hu/Hu^*mice (∼15% of the offspring) relative to the expected 25% Mendelian ratio, indicating ∼40% embryonic lethality in *Stxbp1^Hu/Hu^* mice (**Figure 2E**).

In molecular characterizations from cerebral cortex tissues, *Stxbp1^Hu/Hu^*mice presented only human *STXBP1* mRNA, confirming successful humanization (**Figure 2F**, left and middle panels). Cross-reactive qPCR did not detect significant differences in total *STXBP1* mRNA levels across the different genotypes (**Figure 2F**, right panel). When assessing STXBP1 protein abundance (100% conservation of STXBP1 amino acid sequence between mouse and human), there was a ∼40% reduction in STXBP1 protein levels in *Stxbp1^Hu/Hu^*mice relative to wild-type and *Stxbp1^Hu/+^* littermates (**Figure 2G**). The STXBP1 downregulation observed in *Stxbp1^Hu/Hu^* is not readily explained by a transcriptional mechanism, since total *STXBP1* mRNA levels were unaffected. Post-transcriptional dysregulation of the human *STXBP1* mRNA in the mouse context may contribute to reduced expression, as could the insertion of additional human genomic elements not naturally present in the mouse genome, such as miR-3911 and a lncRNA encoded in the reverse strand (see **Figure 2A**). Such dysregulation would be expected to produce an intermediate reduction in protein expression in *Stxbp1^Hu/+^* mice, which was not observed, raising the possibility that reduced *STXBP1* expression may be a secondary consequence of an unanticipated, pathogenic phenotype resulting from bi-allelic humanization of this locus.

Given that the original *Stxbp1^Hu/Hu^* animals on a pure C57BL/6J genetic background were not viable until they underwent a one-generation outcross with the BALB/c strain, we investigated whether further outcrossing to BALB/c could improve *Stxbp1^Hu/Hu^* viability. After outcrossing to BALB/c for two additional generations, *Stxbp1^Hu/+^* mice with a 3x outcrossed background were subsequently bred together to obtain fully humanized animals, but this unexpectedly increased embryonic lethality (**Supplementary Figure 1**). Finally, we attempted the generation of an *Stxbp1* humanized-disease mouse model by crossing our *Stxbp1^Hu/+^* or *Stxbp1^Hu/Hu^*mice (in 1x and 3x backgrounds) with *Stxbp1* heterozygous knockout mice (*Stxbp1^+/−^*, B6;129S-Stxbp1tm1Sud/J) [8]. Unfortunately, these matings did not produce offspring with *Stxbp1^Hu/-^* genotypes, precluding the generation of an *Stxbp1* humanized haploinsufficiency model.

### *Syngap1* humanized-haploinsufficient mice

To generate a *Syngap1*-disease mouse model that would recapitulate disease-linked phenotypes and allow for testing of human gene-targeted therapies, we crossed the *Syngap1^Hu/Hu^* mice with *Syngap1*-heterozygous mice generated by Kim et. al. (*Syngap1^+/−^*, B6;129-Syngap1^tm1Rlh^/J) in which exons 7 and 8 of *Syngap1* are deleted, resulting in the introduction of a premature termination codon [12] (**Figure 3A**). These matings produced equal ratios (∼50% each) of *Syngap1* humanized-haploinsufficient mice (*Syngap1^Hu/-^*) and hybrid control littermates (*Syngap1^Hu/+^*) (**Figure 3A**), indicating no embryonic lethality. Body weights up to 15 weeks of age showed that *Syngap1^Hu/-^* and *Syngap1^Hu/+^* females were similar in growth rate and size, while *Syngap1^Hu/-^* males were modestly smaller than *Syngap1^Hu/+^*starting at 12 weeks of age (**Figure 3B**).

**Figure 3.**
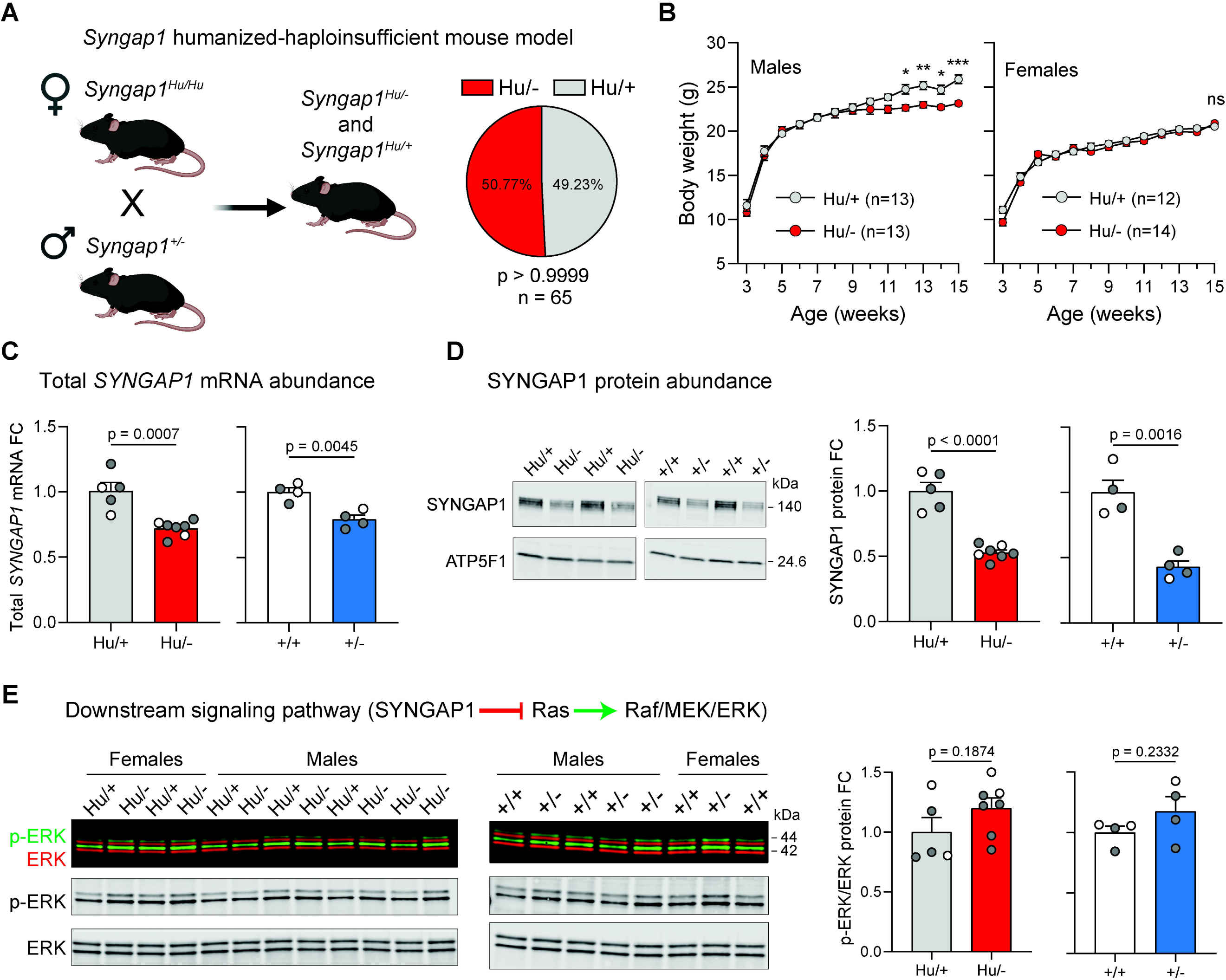
Molecular characterization of a *Syngap1* humanized-haploinsufficient mouse model. (**A**) Cartoon depicting the breeding scheme utilized to generate the *Syngap1* humanized-haploinsufficient mouse model and the corresponding genotypic ratios from the offspring. Created with BioRender. (**B**) Body weights of *Syngap1* humanized-disease mice. (**C**) RT-qPCR from hemi-brain tissue of 34-week-old and 14 to 23-week-old *Syngap1* humanized-haploinsufficient and *Syngap1* heterozygous-haploinsufficient mice, respectively. *Actb* and *Atp5f1* mRNA were used as endogenous control. (**D**) SYNGAP1 western blot from samples in (**C**). ATP5F1 was used as endogenous control. (**E**) Phosphorylated ERK and total ERK western blots from samples in (**C**). (**B**, **C**, **D** and **E**) Data are represented as mean values ± SEM. Data points represent independent biological replicates. (**C**, **D**, **E**) White and gray data points indicate females and males, respectively. (**A**) Binomial test. (**B**) 2-way ANOVA with Sidak’s multiple comparison test (**C**, **D** and **E**) Unpaired t-test. FC, fold change. ns, non-statistically significant. * p < 0.05. ** p < 0.01. *** p < 0.001.

Cross-reactive RT-qPCR of brain RNA extracts from *Syngap1^Hu/-^*revealed a ∼30% reduction in total (mouse and human) *SYNGAP1* mRNA relative to hybrid controls (*Syngap1^Hu/+^*) (**Figure 3C, left**), suggesting that mutant *Syngap1* mRNA is subject to nonsense-mediated decay (NMD) due to the presence of a premature termination codon [for review on NMD, see ref. [26]]. As a control, there was also reduced *Syngap1* mRNA levels (∼20%) in *Syngap1^+/−^* mice relative to wild-types (**Figure 3C, right**), which is in agreement with previous qPCR-based RNA characterizations of this mouse model [11].

Western blotting showed a ∼50% reduction in total SYNGAP1 protein levels in *Syngap1^Hu/-^* hemi-brain tissue, similar to the decrease seen in *Syngap1^+/−^ mice* compared to WT controls (**Figure 3D**). The absence of full SYNGAP1 protein and/or truncated protein species produced by the null allele indicates that the mutant transcripts are either translationally inactive or generate a truncated protein that is readily degraded. Given SYNGAP1’s role as a negative regulator of the Ras-Raf-MEK-ERK signaling pathway [13, 27], ERK signaling was examined by assessing basal levels of phosphorylated ERK (p-ERK) relative to total ERK. *Syngap1^Hu/-^* mice displayed a non-significant trend toward increased p-ERK over total ERK relative to *Syngap1^Hu/+^* (**Figure 3E**), consistent with observations in *Syngap1^+/−^* vs. *Syngap1^+/+^* mice (**Figure 3E**). These data demonstrate the successful generation of a *Syngap1* mouse model that expresses haploinsufficient levels of human SYNGAP1.

### *Syngap1* humanized-haploinsufficient mice exhibit age-dependent behavioral phenotypes

To characterize behavioral phenotypes at multiple ages, juvenile (4 to 5 weeks old) and adult (30 to 31 weeks old) male and female *Syngap1^Hu/+^* and *Syngap1^Hu/-^* underwent a series of locomotor and cognitive tests. 30-minute open field assays showed an increase in total distance traveled (**Figure 4A**) and in stereotypic behaviors (**Figure 4C**) [28] in juvenile *Syngap1^Hu/-^* mice, indicating hyperactivity and cognitive dysfunction in line with prior behavioral characterizations of non-humanized *Syngap1* haploinsufficient mouse models [11, 29]. However, these differences were not preserved in 30-week-old *Syngap1^Hu/-^* mice (**Figure 4B&D**). Neither 4- nor 30-week mice exhibited differences in anxiety-like behaviors in the open field test, as determined by time spent in the center of the arena (**Figure 4E&F**). As other mouse models of SYNGAP1 haploinsufficiency display reduction in anxiety-like behaviors [29–31], elevated plus maze test was also assessed. We observed no differences between *Syngap1^Hu/+^* and *Syngap1^Hu/-^*, irrespective of age, in either the number of open arm entries or time spent in open arms of the maze (**Figure 4G&H**), confirming a lack of an anxiety phenotype in the humanized-haploinsufficient mice. Next, we evaluated locomotor coordination and precision using a horizontal ladder test. While no differences were seen in juvenile mice, 30-week-old *Syngap1^Hu/-^* mice exhibited significantly more foot slips compared to *Syngap1^Hu/+^* mice (**Figure 4I**). Similarly, Muhia et. al. [32] and Nakajima et. al. [31] found deficits in locomotor adaptation in 12- and 53-week-old mice via accelerating rotarod. Therefore, locomotor phenotypes likely represent an age-dependent phenotype and may arise from aberrant plasticity during a period of heightened locomotor plasticity from P30-60 [33]. Finally, we performed the Barnes maze test to assess rodent spatial learning and memory. During the learning phase of the Barnes Maze (first four days), no differences were seen between *Syngap1^Hu/+^* and *Syngap1^Hu/-^* at either age when time to enter the target hole was measured (**Figure 4J**). However, on the fifth day (test day), both juvenile and adult *Syngap1^Hu/-^* mice took significantly longer to contact the target hole, indicating impaired spatial memory and/or diminished memory recall (**Figure 4K**). These findings show that *Syngap1* humanized-haploinsufficient mice recapitulate several behavioral phenotypes associated with *SYNGAP1*-disorder, some of which are age dependent.

**Figure 4.**
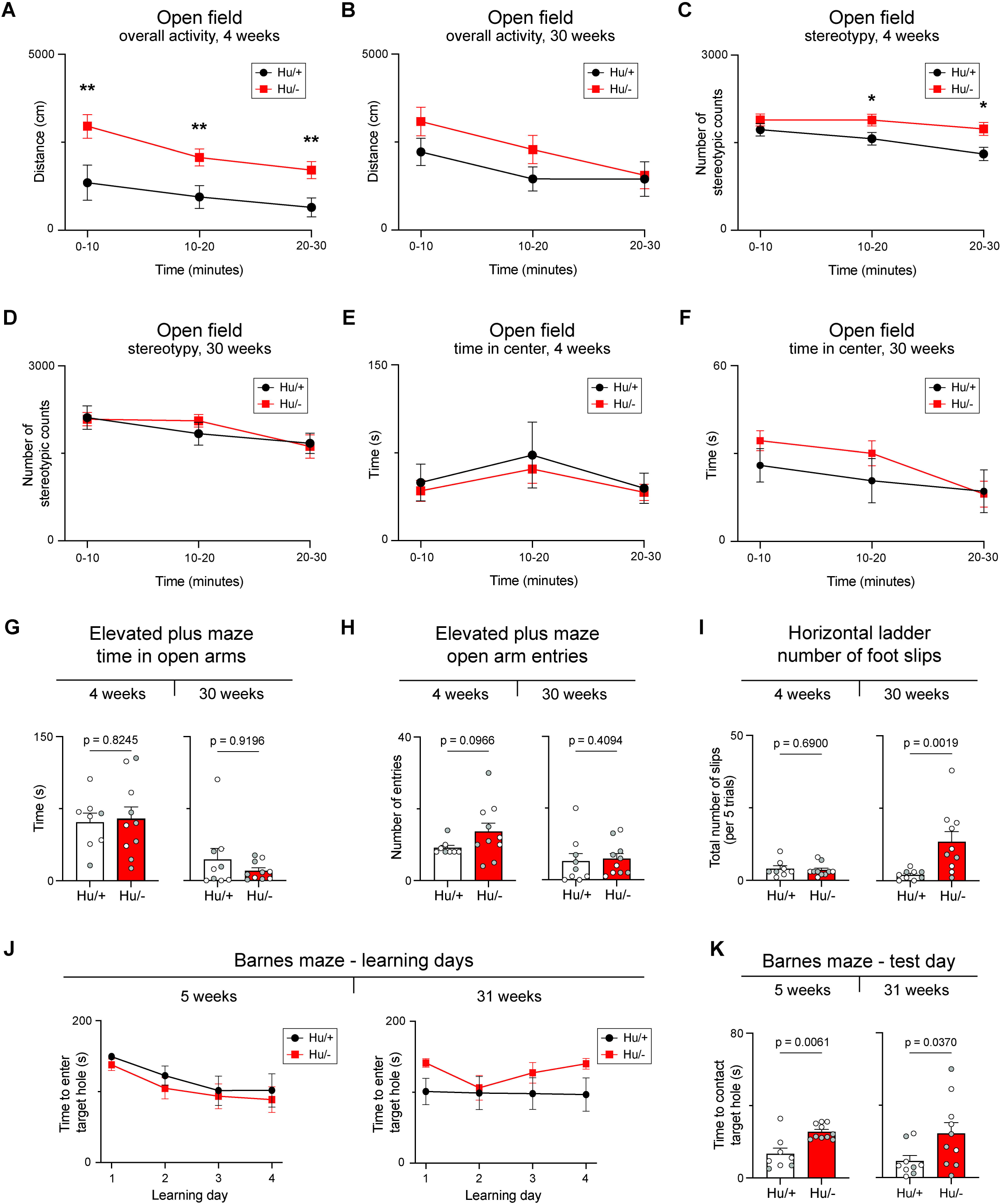
Phenotypic characterization of the *Syngap1* humanized-haploinsufficient mouse model. Open field testing of 4- and 30-week-old mice for (**A** and **B**) overall activity levels via total distance traveled, (**C** and **D**) number of stereotypic behaviors and (**E** and **F**) total time in center of arena (4-week old: Hu/+, *n* = 8; Hu/-, *n* = 10. 30-week old: Hu/+, *n* = 9; Hu/-, *n* = 10). (**G**) Elevated plus maze testing for time in open arms and (**H**) number of open arm entries. (**I**) Horizonal ladder testing from 4- and 30-week-old mice for number foot slips per 5 trials. (**J**) Learning phase of Barnes maze testing from 4- and 30-week-old mice. (**K**) Barnes maze test day, time to find target hole. (**A, B, C, D, E, F**, and **J**) Data are represented by group means ± SEM. (**G, H, I,** and **K**) Data are represented as mean ± SEM. Dots represent individual animals. White and gray data points indicate females and males, respectively. (**A, B, C, D, E, F**, and **J**) Mixed-effects analysis followed by Tukey’s multiple comparisons test when appropriate. Significance stars represent between group comparisons. (**G, H, I,** and **K**) Mann-Whitney test between genotypes. \**p* < 0.05, \*\**p* < 0.01.

### *Syngap1* humanized-haploinsufficient mice display electrophysiological abnormalities

To determine if humanized-haploinsufficient mice exhibit electrophysiological abnormalities, intracranial EEG recordings were obtained from 4 *Syngap1^Hu/+^* and 7 *Syngap1^Hu/-^* mice aged 6 to 9 weeks. Electrodes were implanted into the auditory cortices, barrel cortices, motor cortices, visual cortices, and CA1 of hippocampi bilaterally. Forty-eight hours post-implantation EEGs were recorded continuously for up to 72 hours (**Figure 5A**). To evaluate epileptiform signatures of network excitability, we quantified the number of spike-wave discharges (SWDs) per 12-hour period occurring in at least 4 leads simultaneously. SWDs were detected in all *Syngap1^Hu/-^* mice during both daytime and nighttime but were completely absent in control animals (**Figure 5B**). Power spectral densities in *Syngap1^Hu/-^* mice showed an increase in low-frequency activity as indicated by elevated delta power in all brain regions during both nighttime (**Figure 5C&D**) and daytime (**Supplementary Figure 2A&B**). This robust and widespread generalized slowing likely reflects cerebral dysfunction associated with SYNGAP1-disorder [34–36]. For each brain region, the mean power of major frequency bands was used to calculate alpha-delta (**Figure 5E**) and alpha-theta ratios (**Figure 5F**). *Syngap1^Hu/-^*mice exhibited a significant reduction in alpha-delta ratio in the motor cortex during both daytime and nighttime. In the hippocampus, alpha-theta ratios showed a trend toward reduction at both times. Differences in normalized band power between genotypes were only observed in the auditory cortex during the daytime (**Supplementary Figure 2C-G**), while spike frequency differed only in the hippocampus at nighttime (**Supplementary Figure 2H&I**). Overall, these findings demonstrate that *Syngap1^Hu/-^* mice display distinct EEG features that correspond with epileptiform activity and generalized background slowing.

**Figure 5.**
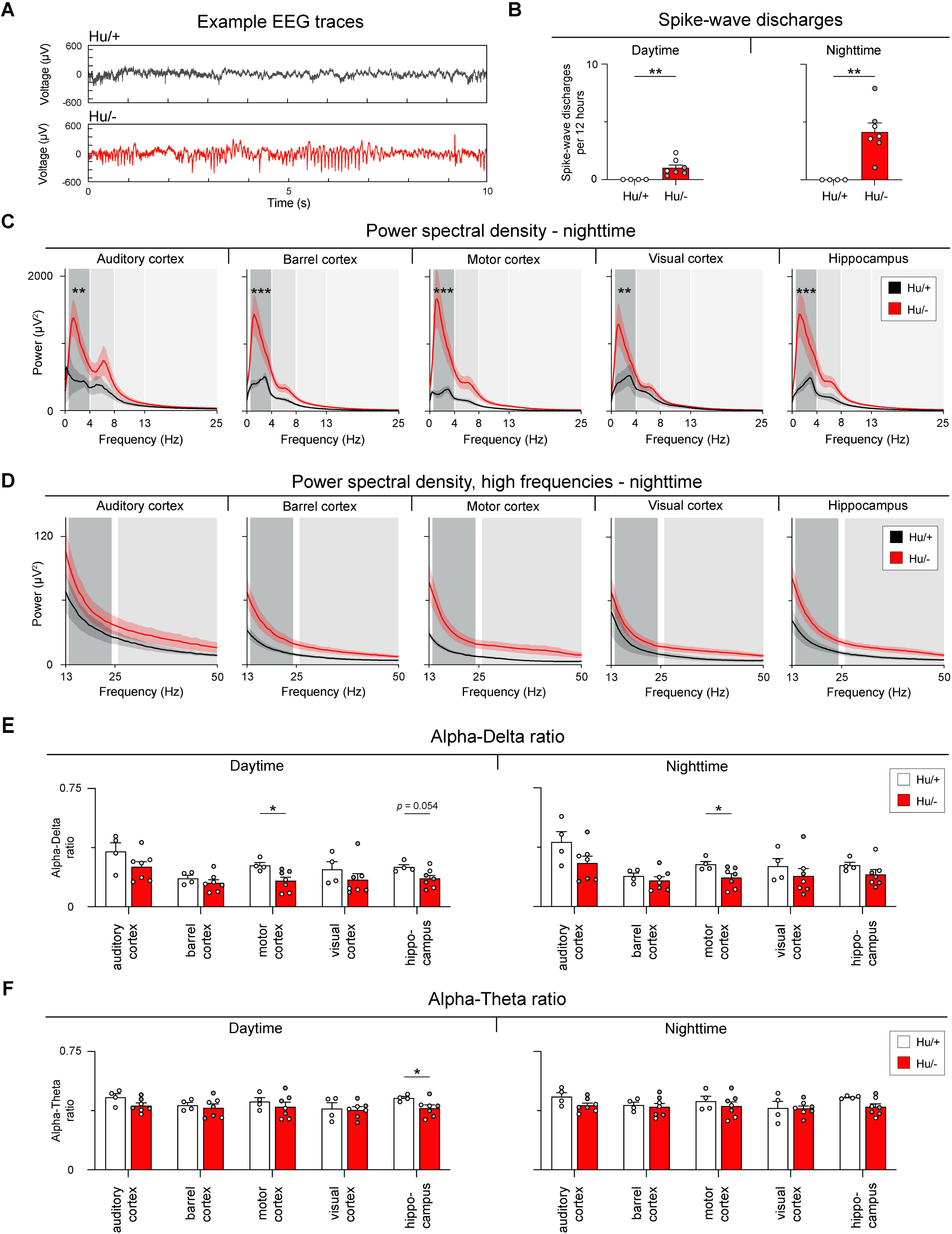
EEG mapping of the *Syngap1* humanized-haploinsufficient mouse model. (**A**) Representative EEG traces from Hu/+ and Hu/- mice. (**B**) Number of spike-wave discharges per 12 hours for daytime and nighttime (Hu/+, *n* = 4; Hu/-, *n* = 7). (**C**) Nighttime low frequency and (**D**) high frequency power spectral densities by brain region. Shaded gray backgrounds represent EEG frequency bands. Delta 0-4Hz, Theta 4-8Hz, Alpha 8-13Hz, Beta 13-25Hz, and Gamma 25-50-Hz. (**E**) Alpha-delta ratios by brain region for daytime and nighttime. (**F**) Alpha-theta ratio by brain region for daytime and nighttime. (**B, E,** and **F**) Data are represented as mean ± SEM. Dots represent individual animals, white and gray data points indicate females and males, respectively. (**C** and **D**) Lines represent group means and shading represents ± SEM. (**B**) Mann-Whitney. (**C** and **D**) Linear mixed model to compare area under the curve for each power spectra within each brain region as a function of strain, frequency, and the interaction between strain and frequency. Post hoc comparisons were corrected using Tukey’s HSD. (**E** and **F**) Unpaired t test within regions. \*\**p* < 0.01, \*\*\**p* < 0.001.

### EEG analysis of individuals with *SYNGAP1*-disorder reveal spike-wave discharges and low-frequency shifts

To evaluate whether individuals with *SYNGAP1*-disorder show electrophysiological signatures that parallel our findings in *Syngap1^Hu/-^*mice, we analyzed surface EEGs from control and *SYNGAP1*-disorder participants (**Figure 6A**). The presence and frequency of SWDs were quantified from SmartForms for 19 participants (52 EEGs total). SWDs were present in 63% of participants and 40.4% of EEGs (**Figure 6B**). For power spectral densities comparisons, age-matched groups - control (84 EEGs, *n* = 84) and *SYNGAP1*-disorder (21 EEGs, *n* = 21) – were analyzed and compared (**Supplementary Figure 3**). We observed regional increases in low-frequency power in parietal and central leads (**Figure 6C&D**). *SYNGAP1* participants also showed decreased alpha-delta ratios in parietal and occipital leads (**Figure 6E**) and, as predicted by Galer et. al. [37], a brain-wide reduction in alpha-theta ratios (**Figure 6F**). To assess age differences, participant data were stratified into younger (≤10 years of age) and older (> 10 years of age) groups (**Supplementary Figures 4A&B and 5A&B, respectively**). The low-frequency shift was also seen in parietal and central leads in younger *SYNGAP1* participants (**Supplementary Figure 4C-D**) and extended to frontal and temporal leads in older participants (**Supplementary Figure 5C-D**). Similar decreases in alpha-delta and alpha-theta ratios were present in both age groups (**Supplementary Figures 4E&F and 5E&F**). Overall, these human EEG signatures are parallelled by those seen in the *Syngap1* humanized-haploinsufficient model and provide translational biomarkers to assess therapeutic efficacy and predict the potential clinical effect of human gene-targeted therapies.

**Figure 6.**
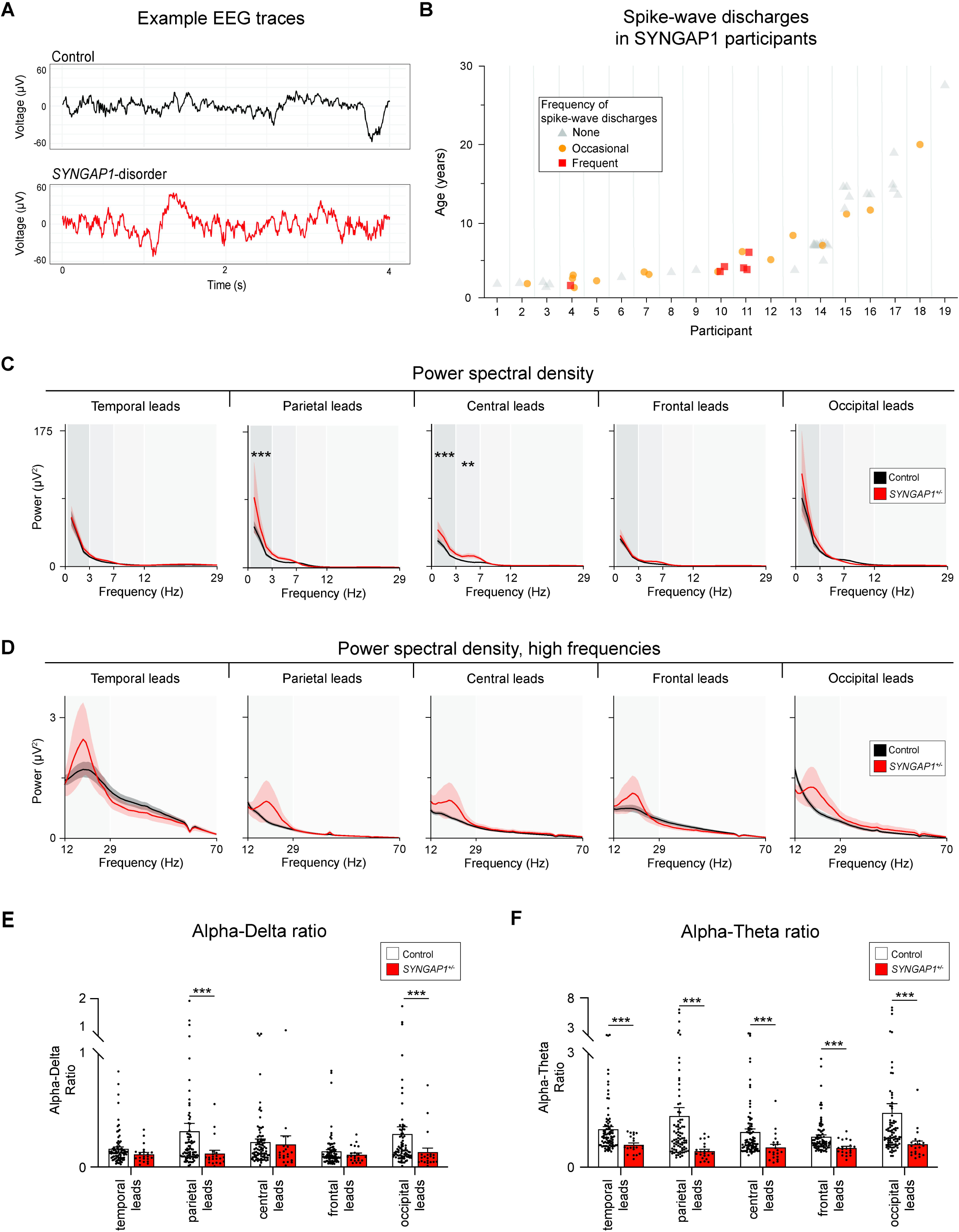
Quantitative EEG analysis in age-matched *SYNGAP1* participants. **(A)** Representative EEG traces from a control and a *SYNGAP1*-disorder participant. (**B**) Presence and frequency of spike-wave discharges in *SYNGAP1*-disorder participants (participants, *n* = 19; EEGs, *n* = 52). Individual shapes represent 1 EEG recording and shape/color represent frequency of spike-wave discharges from SmartForms. (**C**) Low frequency and (**D**) high frequency power spectral densities by leads/region (control individuals, *n* = 84; *SYNGAP1*-disorder participants, *n* = 21. 1 EEG from each individual was included.). (**E**) Alpha-delta ratio by leads/region (**F**) Alpha-theta ratio by leads/region. (**C** and **D**) Lines represent group means and shading represents ± SEM. (**E** and **F**) Bars represent group means ± SEM. (**C** and **D**) Linear mixed model to compare area under the curve for each power spectra within each brain region as a function of group, frequency, and the interaction between group and frequency. (**E** and **F**) Mann-Whitney test within regions.\*\**p* < 0.01, \*\*\**p* < 0.001.

### Human *SYNGAP1*-targeted ASOs modulate *SYNGAP1* expression in *Syngap1^Hu/-^* neurons

Our final goal was to determine whether human *SYNGAP1* expression could be modulated in the *Syngap1* humanized-haploinsufficient mouse model using a human-specific *SYNGAP1* therapy. We and others had previously identified an alternative 3’ splice site (Alt. 3’ss) event in *SYNGAP1* exon 10-11 that promotes NMD and restricts *SYNGAP1* expression (**Figure 7A**) [16, 17]. This Alt. 3’ss event is dependent on the binding of polypyrimidine tract binding proteins (PTBPs) to *SYNGAP1* pre-mRNA, opening the opportunity for a therapeutic splice switching strategy using steric-blocking ASOs to specifically disrupt PTBP binding and upregulate *SYNGAP1* (**Figure 7A**). Our previous study identified two main ASO candidates named ET-019 and ET-085 targeting PTBP binding sites located upstream (Site-1) and downstream (Site-2) from the Alt. 3’ss, respectively, that redirected splicing and increased *SYNGAP1* mRNA abundance in *SYNGAP1* patient-derived iPSC-neurons [17]. Here we tested these human-specific compounds in *Syngap1^Hu/-^* derived primary neurons to examine their ability to upregulate human *SYNGAP1*. Gymnotic delivery of ET-019 and ET-085 in *Syngap1^Hu/-^* -derived primary neurons was well tolerated and significantly increased *SYNGAP1* mRNA abundance (1.7- and 1.5-fold, respectively) relative to mock-treated cells or a negative control ASO (ET-SC2) with the same length and chemistry (**Figure 7B**). In follow-up ASO testing, we found that ET-019 upregulated human *SYNGAP1* in a dose-dependent fashion, reaching a ∼2.9-fold increase at 20 µM concentration (**Figure 7C**), indicating effective human target engagement and improved human *SYNGAP1* expression in a haploinsufficient mouse model. In parallel, we also investigated whether *SYNGAP1* expression could be downregulated using a gapmer ASO (Gap-SYN) specifically designed to trigger RNase H1-mediated degradation of human *SYNGAP1* mRNA. *Syngap1^Hu/-^* primary neurons treated with Gap-SYN showed a strong reduction (∼70%) in *SYNGAP1* mRNA levels, confirming successful knockdown of the target (**Figure 7B**). Together, our data indicates that *SYNGAP1* expression can be modulated bidirectionally in the humanized haploinsufficiency context using human gene-targeted ASOs.

**Figure 7.**
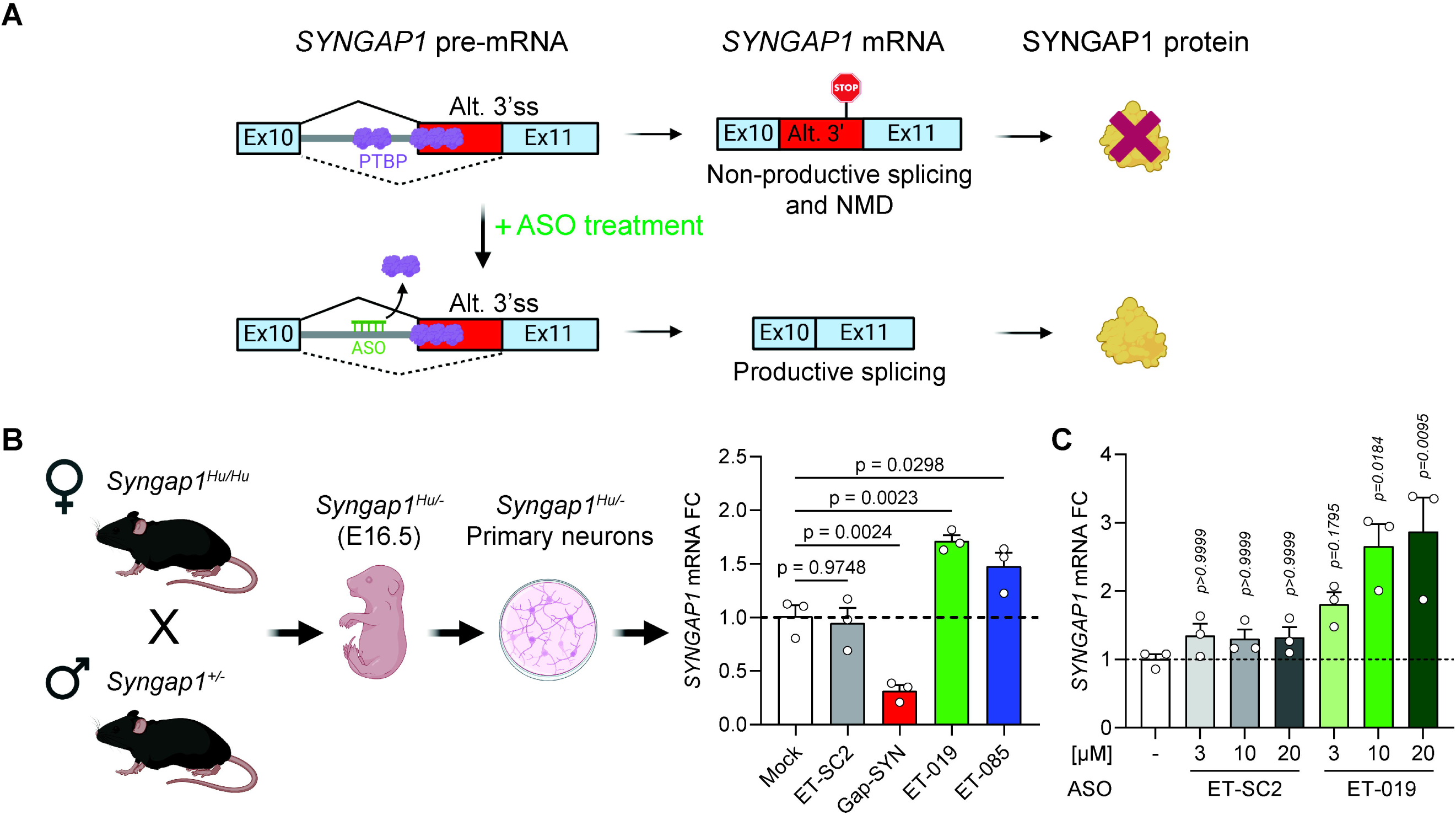
Bidirectional modulation of human *SYNGAP1* expression in *Syngap1^Hu/-^*-derived primary neurons using ASOs. (**A**) Cartoon schematic representing the ASO-mediated splice-switching approach to upregulate human *SYNGAP1* expression by disrupting PTBP binding to *SYNGAP1* pre-mRNA. (**B**) Left, cartoon depicting the breeding scheme used to obtain *Syngap1^Hu/-^* embryos for primary neuron isolation. Created with Biorender. Right, RT-qPCR from *Syngap1^Hu/-^* primary cortical neurons treated with ASOs for 7 days. All ASOs were added at 10 µM except for Gap-SYN, which was added at 3 µM. ET-SC2 was used as a non-targeting ASO control. (**C**) RT-qPCR from *Syngap1^Hu/-^* primary cortical neurons treated with ASOs for 7 days. *Atp5f1* mRNA was used as endogenous control. A qPCR assay spanning exons 16-17 of human *SYNGAP1* was used. *Atp5f1* mRNA was used as endogenous control. (**B**) One-way ANOVA with Dunnett’s multiple comparison test vs. mock-treated cells (−). (**C**) Kruskal-Wallis with Dunn’s multiple comparison test vs. mock-treated cells. Data are represented as mean values ± SEM. Data points represent independent biological replicates. Alt. 3’ss, alternative 3’ splice site. FC, fold change.

## Discussion

Here we generated humanized *Syngap1* and *Stxbp1* mouse models to enable human gene-targeted therapeutic development for *SYNGAP1* and *STXBP1* disorders. We further demonstrate construct validity for a *Syngap1* humanized-disease model (*Syngap1^Hu/-^*), which recapitulates haploinsufficiency, hyperactivity, cognitive, motor, and electrophysiological abnormalities observed in patients with *SYNGAP1*-disorder.

*Stxbp1^Hu/Hu^* mice were non-viable until outcrossed one generation to BALB/c. While the genetic cause for the impaired viability of *Stxbp1^Hu/Hu^* mice is unclear, further outcrossing to BALB/c did not ameliorate embryonic lethality, and precluded the generation of *Stxbp1* humanized-disease mice (*Stxbp1^Hu/-^*). However, hybrid *Stxbp1^Hu/+^* mice were viable and expressed normal levels of STXBP1, providing a useful model to assess human target engagement and mechanistic interrogations of clinical drug candidates.

Our *Syngap1* humanization also included the human *SYNGAP1* antisense transcript (*SYNGAP1-AS*), which may modulate *SYNGAP1* expression and as such presents a potential therapeutic target available only in this line. While all *Syngap1^Hu/-^* animals have consistent SYNGAP1 loss-of-function, a weak inverse correlation between SYNGAP1 and active ERK levels, as indicated by ERK phosphorylation, was observed. Alternative cellular phenotypes such as AMPA receptor trafficking or dendritic spine morphology [38] may represent more sensitive measures to detect downstream rescue of human SYNGAP1 haploinsufficiency.

Consistent with other *Syngap1^+/−^* mouse lines, *Syngap1^Hu/-^*mice displayed disease-related behaviors including hyperactivity, increased repetitive behaviors, and memory deficits [12]. However, in aged mice, activity levels normalized and deficits in locomotor coordination and adaptation developed (**Figure 4**), suggesting impaired motor commands and/or altered sensorimotor feedback during ongoing movement. Diverging from other *Syngap1^+/−^* mouse models, *Syngap1^Hu/-^*mice showed no differences in anxiety-like behaviors [14, 15, 31]. The lack of anxiety-related behaviors in multiple tests may reflect differences in human *SYNGAP1* transcript stability or translation efficiency compared to mouse transcripts, leading to a slight increase in protein production beyond that of endogenous mouse *Syngap1* alleles (**Figure 1**). Additionally, we cannot rule out the potential effect of genetic background as it has impacted behavioral phenotypes in other neurodevelopmental disease models [3, 39].

EEG analyses in *Syngap1^Hu/-^* mice and in individuals with *SYNGAP1*-disorder reinforce the face validity of this new mouse model by revealing convergent electrophysiological phenotypes. Spike-wave discharges - indicative of absence seizures [23, 40] – occurred in *Syngap1^Hu/-^* mice and were present in 12 of 19 SYNGAP1 participants or 40.4% of annotated EEGs. While *Syngap1^Hu/-^* mice displayed an elevated absolute power spectral density, SYNGAP1 patients exhibited increased power confined to low frequency bands (i.e. delta and theta). Generalized slowing of EEG rhythm was brain-wide in mice but restricted to parietal and central leads in participants. However, generalized slowing was further confirmed in participants by regional reductions in alpha-delta ratios and a broad decrease in alpha-theta ratios (**Figure 5**). These low-frequency EEG shifts are not unique to *SYNGAP1*-disorder as similar patterns are reported in multiple encephalopathies [41, 42], and in other CNS disorders including temporal lobe epilepsy [43], ischemic stroke [44], chronic pain, Parkinson’s disease [45], and Angelman syndrome [46]. In Angelman syndrome, elevated delta power predicts lower performance on the Bayley Scale of Infant and Toddler Development and correlates with reduced cognitive, motor, and language performance. Whether similar relationships are seen in *SYNGAP1*-disorder will be determined by the ongoing ProMMiS natural-history study (NCT06555965). Given the presence of disease-relevant behaviors and EEG slowing found here in *Syngap1^Hu/^*^−^ mice, we anticipate the ProMMiS study will reveal similar associations between EEG slowing and functional outcomes [47].

EEG signals predominately reflect synchronized synaptic currents [48, 49] with low-frequency rhythms arising from thalamic and cortical generators [50–53]. Accordingly, the low-frequency shift observed here may reflect weakened thalamocortical input and/or altered inhibitory interneuron function. Generalized slowing is linked to thalamocortical deafferentation in numerous conditions [45, 54, 55], and consistent with this, *Syngap1^+/−^*mice show weakened thalamocortical inputs onto layer-5 pyramidal neurons in the primary sensory cortex [56]. This suggests that thalamocortical deafferentation likely contributes to the low-frequency shift in *SYNGAP1*-disorder. Additionally, impaired maturation and altered synaptic drive of parvalbumin (PV) interneuron in *Syngap1^+/−^* mice [57] may further bias these networks to slower rhythms, as PV mediated inhibition typically suppresses low frequencies and supports faster oscillations [58, 59]. These mechanisms are not mutually exclusive as perineuronal nets modulate thalamocortical inputs to cortical PV neurons [60], and together likely contribute to the low-frequency shifts seen here in both mice and participants.

Finally, we show that two human-specific ASOs [17] can effectively modulate human *SYNGAP1* expression in *Syngap1^Hu/-^* primary neurons (**Figure 7B&C**). As these splice-switching oligos act through PTBP disruption [17], our data suggest that mouse PTBP proteins can also recognize and regulate the Alt. 3’ss of human *SYNGAP1*. Ultimately, the *Syngap1^Hu/-^* mouse model generated here will enable *in vivo* testing of human gene-targeted therapies, and the conserved electrophysiological signatures among SYNGAP1 patients and mice provide translational biomarkers to accelerate the arrival of these therapies to the clinic.

## Author contributions

A.J.F., B.L.B., M.J.B., B.L.D and B.L.P. contributed to experimental design. A.J.F., B.L.B., I.H.O, T.W., R.R., M.J.G., N.M., J.D.M., D.R. performed the experiments and collected the data. A.J.F., B.L.B., K.U. maintained and expanded the mouse colonies. A.J.F., B.L.B., M.H., M.J.G., I.M., J.L.M. performed data analysis. Writing original draft: A.J.F., B.L.B. Writing - Review & Editing: A.J.F., B.L.B, J.L.M., I.M, B.L.P., B.L.D. Funding acquisition: J.L.M., I.H., B.L.P, B.L.D. All authors approved the final manuscript.

## Acknowledgements

This work was supported by R21 NS118280 from the National Institute of Neurological Disorders and Stroke (NIH-NINDS) and by a sponsored research agreement with Ionis Pharmaceuticals to B.L.P. and B.L.D., by the Neurodevelopmental Disabilities T32 Training Grant T32NS007413 from NIH-NINDS to B.L.B, The Center for Epilepsy and Neurodevelopmental Disorders (ENDD), R01 NS127830-01A1 and R01 NS131512-01 from NIH-NINDS to I.H., K23 NS140491-01A1 from NIH-NINDS to J.L.M., and the American Academy of Neurology (AAN), American Epilepsy Society (AES), the Epilepsy Foundation, & the American Brain Foundation (ABF) through the Susan Spencer Award to J.L.M. We thank Dr. Elizabeth A. Heller (University of Pennsylvania) for assisting with the design of the humanized mouse models and Phillip Morrin (Children’s Hospital of Philadelphia) for assisting with mouse husbandry.

