## Supplementary Information for "Translatable electrophysiological and behavioral abnormalities in a humanized model of *SYNGAP1*-disorder"

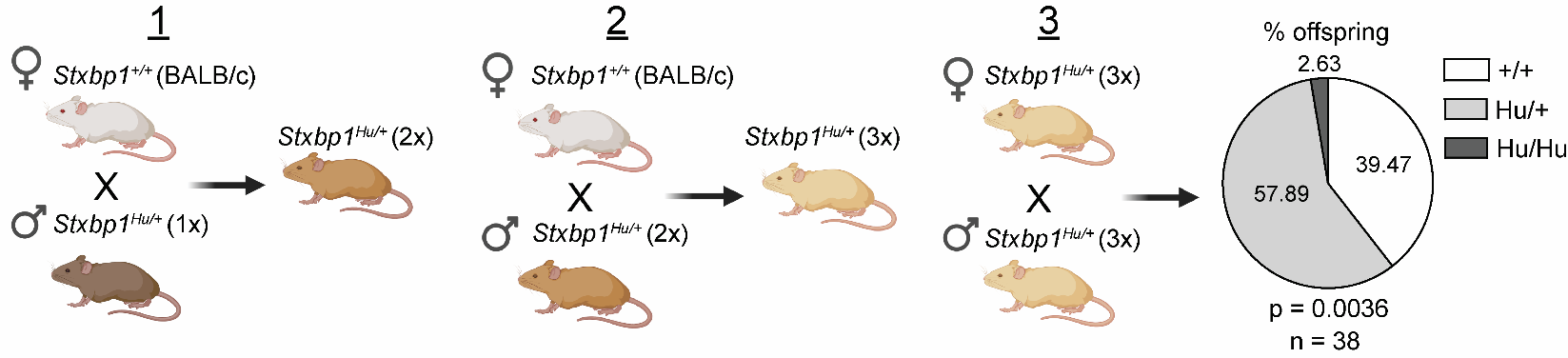
**Supplementary Figure 1. Outcrossing the *Stxbp1* humanized mice to the BALB/c strain.** Cartoon depicting the breeding scheme utilized to outcross the *Stxbp1* humanized mouse model to the BALB/c strain for two additional generations as well as the corresponding genotypic ratios from the offspring of *Stxbp1^Hu/+^* x *Stxbp1^Hu/+^* matings in the 3x outcrossed background. Created with BioRender. Chi-square test (df = 2, *n* = 38).

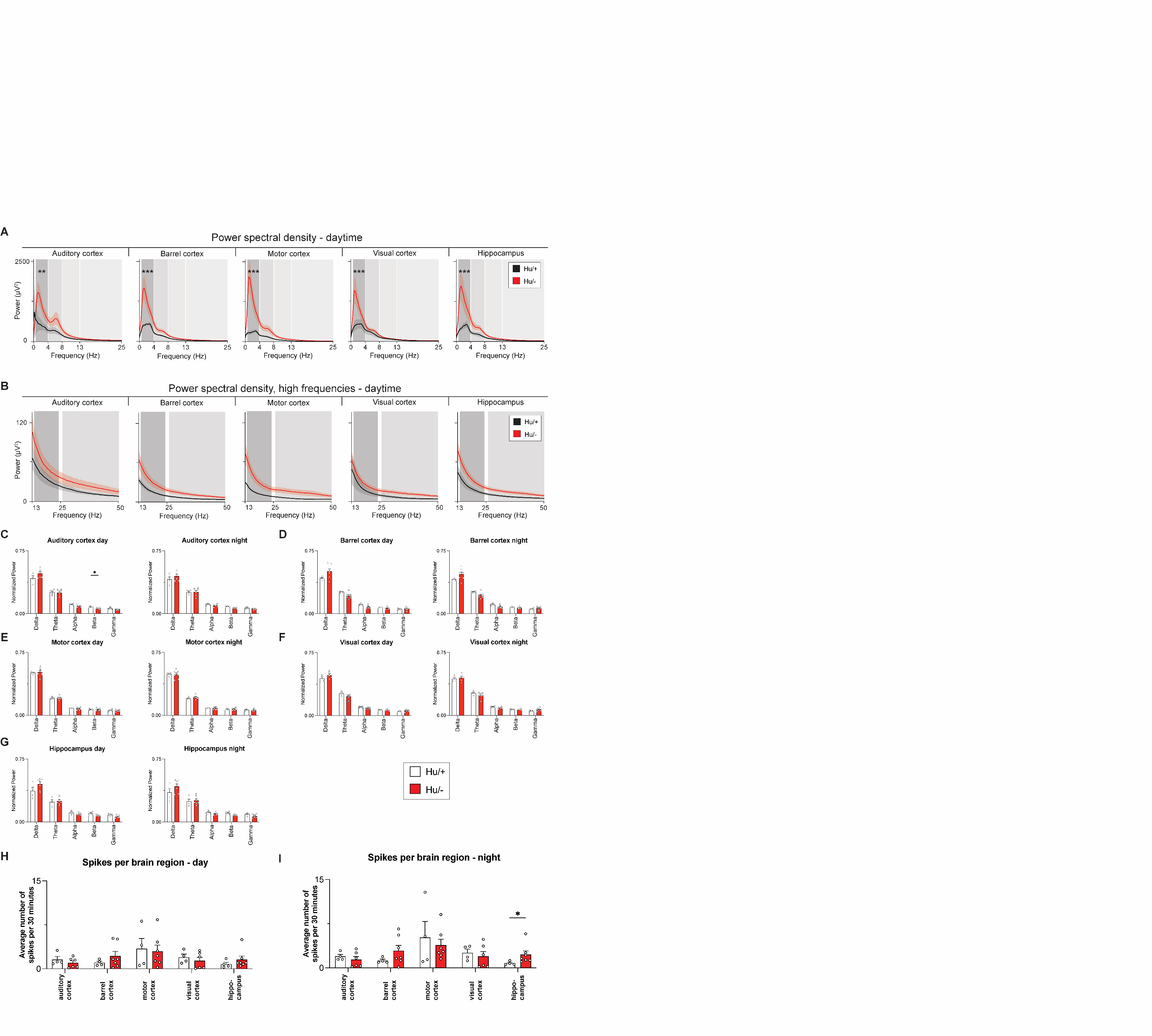

**Supplementary Figure 2. Additional EEG measures from *Syngap1* humanized-haploinsufficient mice.** Normalized band powers from both daytime and nighttime for the (**A**) auditory cortex, (**B**) barrel cortex, (**C**) motor cortex, (**D**) visual cortex, and (**E**) hippocampus (Hu/+, *n* = 4; Hu/-, *n* = 7). Average number of spikes per 30-minute period for each brain region during (**F**) daytime and (**G**) nighttime. Data are represented as mean ± SEM. Dots represent individual animals. (**A-E**) Individual Mann-Whitney tests per region. (**C** and **D**) Mixed-effects model. Fixed effects for frequency, genotype, and frequency x genotype. Sidak’s multiple comparisons test was used when appropriate. **p* < 0.05.

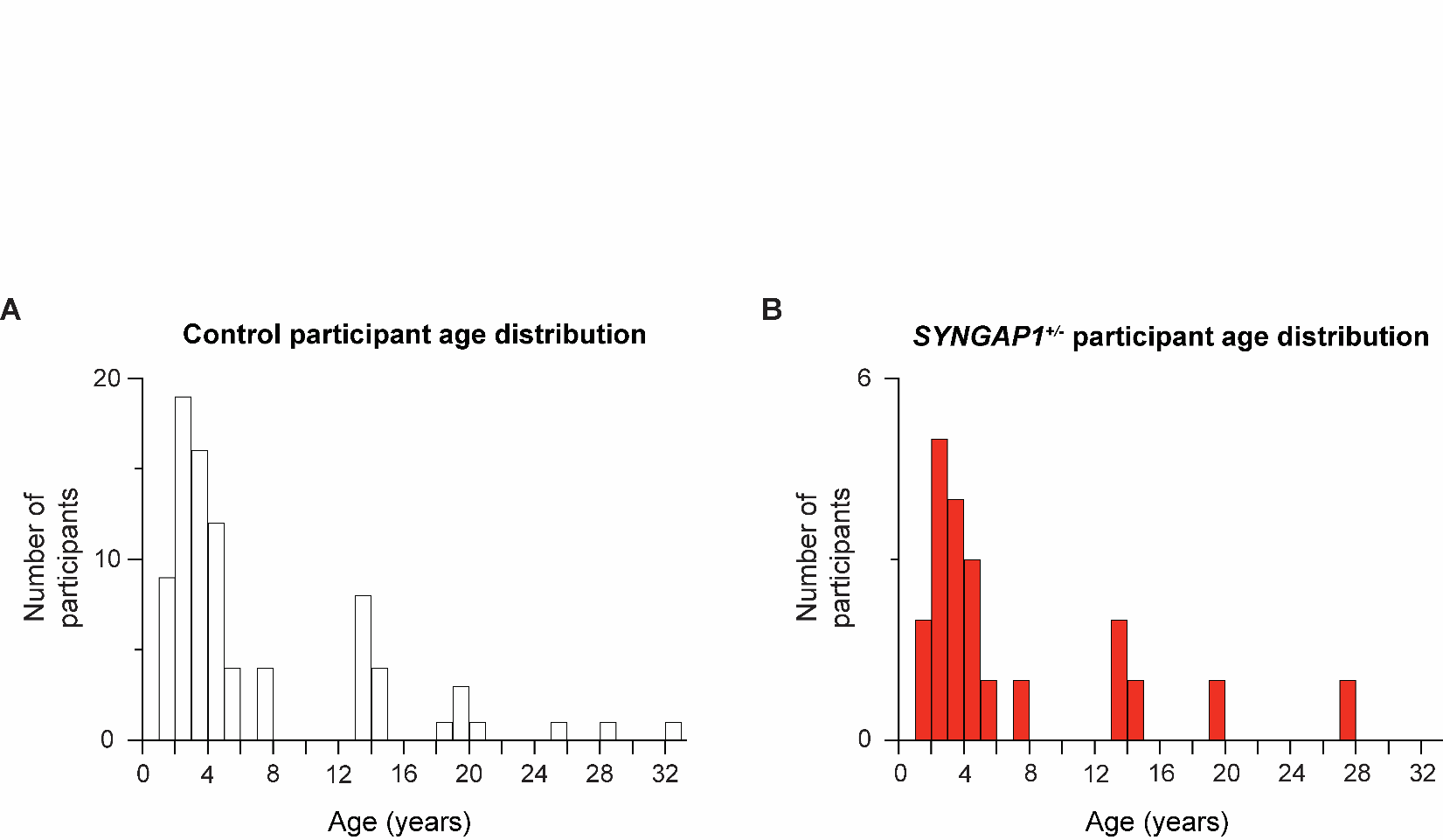

**Supplementary Figure 3.** Age distributions of (**A**) control and (**B**) *SYNGAP1*-disorder participants that were analyzed in Figure 6.

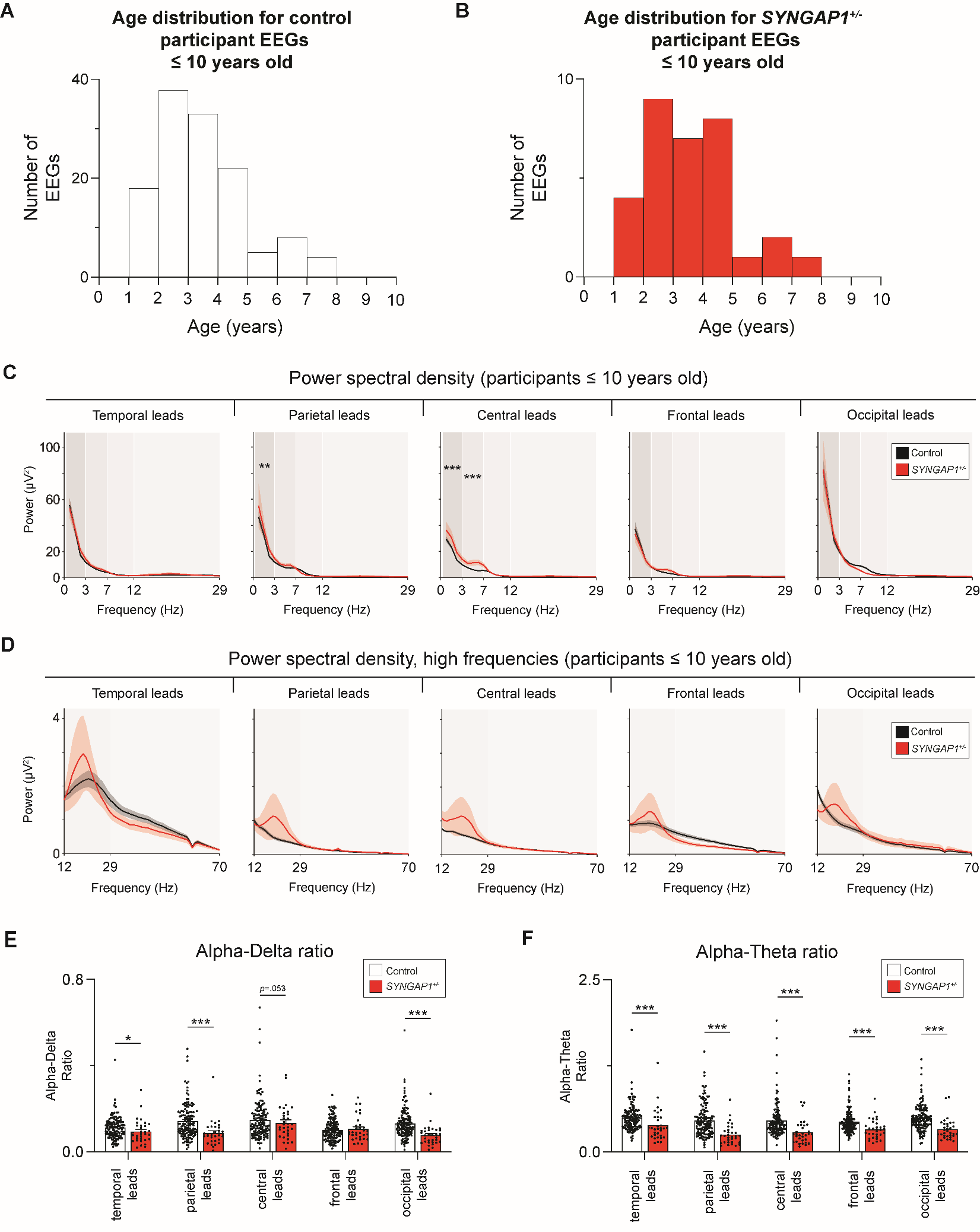

**Supplementary Figure 4. EEG analysis in younger (≤10 year) *SYNGAP1* participants.** Age distributions of (**A**) control and (**B**) *SYNGAP1*-disorder participants (control individuals, *n* = 128 EEGs; *SYNGAP1* participants, *n* = 32 EEGs). (**C**) Low frequency and (**D**) high frequency power spectral densities by leads/region. (**E**) Alpha-delta ratio by leads/region. (**F**) Alpha-theta ratio by leads/region. (**C** and **D**) Lines represent group means and shading represents ± SEM. (**E** and **F**) Bars represent group means ± SEM. (**C** and **D**) Linear mixed model to compare area under the curve for each power spectra within each brain region as a function of group, frequency, and the interaction between group and frequency. (**E** and **F**) Mann-Whitney test within regions. **p*<0.05, ***p* < 0.01, ****p* < 0.001.

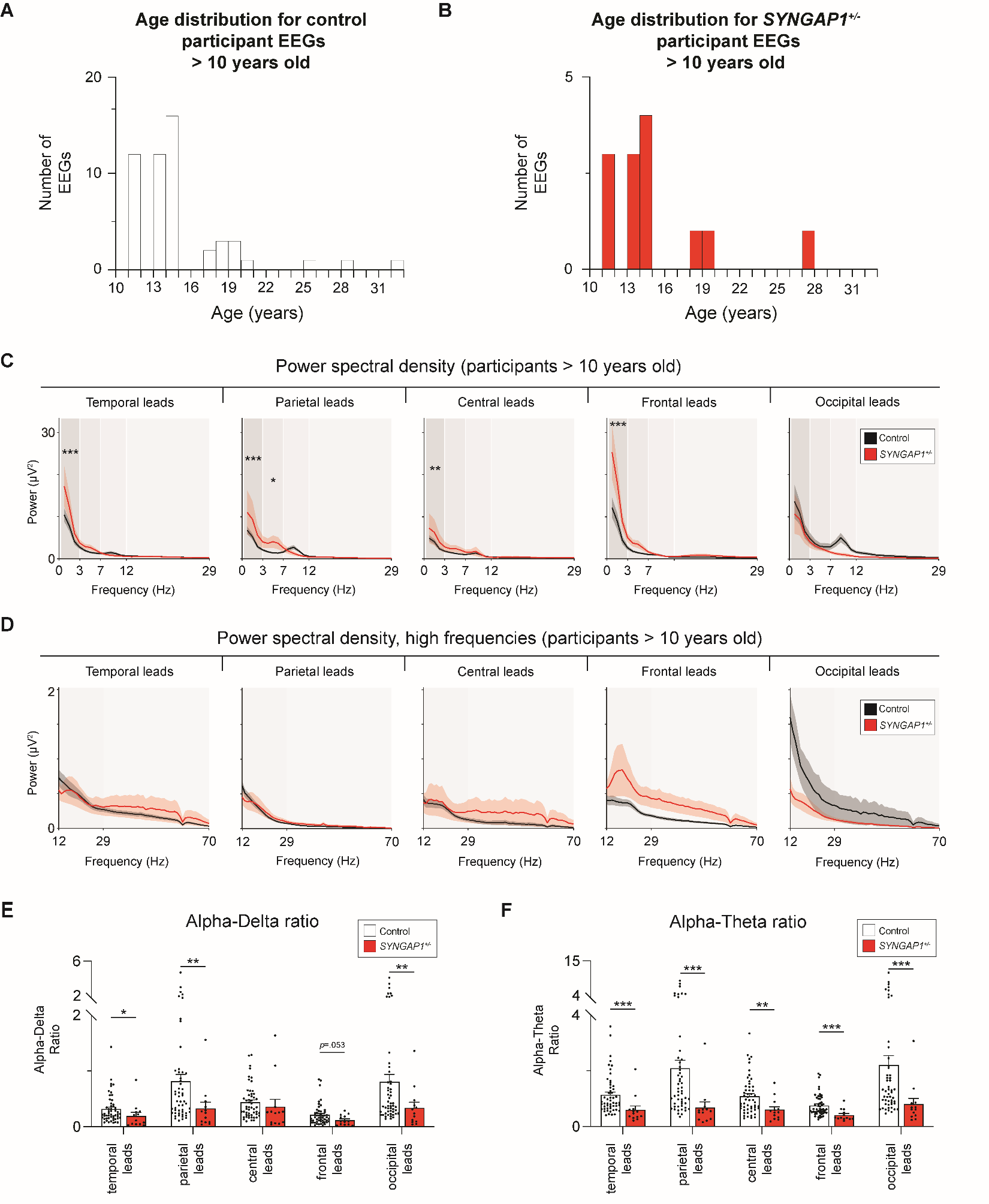

**Supplementary Figure 5. EEG analysis in older (>10 year) *SYNGAP1* participants.** Age distributions of (**A**) control and (**B**) *SYNGAP1*-disorder participants (control individuals, *n* = 52 EEGs; *SYNGAP1* participants, *n* = 13 EEGs). (**C**) Low frequency and (**D**) high frequency power spectral densities by leads/region. (**E**) Alpha-delta ratio by leads/region (**F**) Alpha-theta ratio by leads/region. (**C** and **D**) Lines represent group means and shading represents ± SEM. (**E** and **F**) Bars represent group means ± SEM. (**C** and **D**) Linear mixed model to compare area under the curve for each power spectra within each brain region as a function of group, frequency, and the interaction between group and frequency. (**E** and **F**) Mann-Whitney test within regions. **p*<0.05, ***p* < 0.01, ****p* < 0.001.

**Supplementary Table 1. qPCR assays used in this study.**

| **Copy number variation/Genotyping assay for *Syngap1* and *Stxbp1* humanized mouse models** | | | | |
| --- | --- | --- | --- | --- |
| **Name** | **Forward (5’->3’)** | **Reverse (5’->3’)** | **Probe (5’->3’)** | **Assay ID (if applicable)** |
| Mouse_Syngap1_gDNA | GCAGTTGTGTGTTCATCTGTTC | CAGCCCTCATTCACCTCTTT | TGAGCATCAGTACGGGCAAAGCAT | NA |
| Human_SYNGAP1_gDNA | CCCTGACCTCTCTTCTGATAG | CTCTGGCCTAGGGTAAATG | CTGTTCCTGTCCAACCATCACTG | NA |
| Mouse_Stxbp1_gDNA | TCAAGAGAGTTGGTGACAGA | CCCTTTGCCCCTTCAGTTTC | TTTGAAAGGAATTCTAGGGGATCG | NA |
| Human_STXBP1_gDNA | CAGGATCTCAGTGTGAAGCTAAG | GGACAGGGCTTACAATCCTAAA | TCTGGTTTGGTGTGACAGGCTCAG | NA |
| Mouse_Tert_gDNA | GAGACAATGGGTGGCAGTAA | GCTTGGAGTCAGAGACCATAAG | ATGCAGTCCGTGGTTGGATGAGTT | NA |
| **Genotyping assays to detect the presence of *Syngap1* or *Stxbp1* null alleles** | | | | |
| Name | Forward (5’->3’) | Reverse (5’->3’) | Probe (5’->3’) | Assay ID (if applicable) |
| Mouse_Stxbp1_WT_gDNA | CCCTGAGCTCTCCTCTTCCTT | GCAGCAGGAGGACAGCAT | CCACCACCAGCACCTG | NA |
| Mouse_Stxbp1_null_gDNA | GCTTATCGAGCTTGATATCGAATTCCT | CTCAAGTTCAGAATTTGCCTACTTCACC | CAGCCGTGTCACACAC | NA |
| Mouse_Syngap1_WT_gDNA | ACCTCAAATCCACACTCCTCTCCAG | AGGGAACATAAGTCTTGGCTCTGTC | Not applicable | NA |
| Mouse_Syngap1_null_gDNA | ATGCTCCAGACTGCCTTGGGAAAAG | AGGGAACATAAGTCTTGGCTCTGTC | Not applicable | NA |
| **RT-qPCR assays for molecular characterizations** | | | | |
| Name | Forward (5’->3’) | Reverse (5’->3’) | Probe (5’->3’) | Assay ID (if applicable) |
| Mouse_Syngap1_mRNA | GAGTGAGAAGCGCTTGAGA | TTCTTGGGCTCAGGCAG | CAGCAGCAGGTGGAGAAGGACT | NA |
| Human_SYNGAP1_mRNA | GCAGAGTGAGAAGAGGCTA | TCCTCCACCAGCATCAG | TCCCAGATCAAGAGCATCATTGGCA | NA |
| Total_SYNGAP1_mRNA | GCTGGATGAGGATGAGATACAC | GAGCGACCCAAGTGGTATT | AACAAACTGCTGAGACGCAC | NA |
| Human_SYNGAP1_Ex16-17 | Proprietary | Proprietary | Proprietary | IDT (Hs.PT.58.4622325) |
| Mouse_Stxbp1_mRNA | CGTATCAGTGAGCAGACCTA | GGTAGTGCTTTGTATCCAGC | AGACATTATGGAGGACACTATCGAAGACA | NA |
| Human_STXBP1_mRNA | CATCAGCGAGCAGACCTA | GGTAGTGTTTGGTGTCAAGT | AGGACATCATGGAGGACACTATTGAGGA | NA |
| Total_STXBP1_mRNA | ACAAGCACATCGCAGAGG | TTCTTCAGCATCTGGGACAG | AGGAAGTCACCCGGTCTCTGAA | NA |
| Mouse_Atp5f1_mRNA | Proprietary | Proprietary | Proprietary | Thermo (Mm05814774_g1) |
| Mouse_Actinb_mRNA | Proprietary | Proprietary | Proprietary | Thermo (Mm02619580_g1) |

**Supplementary Table 2. ASOs used in this study.**

| **Target** | **ID** | **ASO (5'->3')** | **Length (nt)** | **Chemistry** | **Comments** |
| --- | --- | --- | --- | --- | --- |
| Non-targeting control | ET-SC2 | AGGTCGGTGACAGTTGCA | 18 | 2'-Methoxyethyl (2'MOE) with full phosphorothioate (PS) backbone |  |
| Human SYNGAP1 | Gap-SYN | CGGTGTTTCGGAACTCGC | 18 | Locked nucleic acid (LNA)-DNA-LNA (4-10-4). All PS. |  |
| Human SYNGAP1 | ET-019 | CACGTGGGAGAGAGATGG | 18 | 2'MOE-PS | From McKenna and Felix et. al. 2023 |
| Human SYNGAP1 | ET-085 | TCCAGGGAACATGCTGAG | 18 | 2'MOE-PS | From McKenna and Felix et. al. 2023 |
